# *qg*: Configuration-Driven, Multi-Vendor Acquisition Queue Generation with Reproducible Run-Order and QC Control for Mass Spectrometry

**DOI:** 10.64898/2026.07.03.736300

**Authors:** Witold E. Wolski, Leonardo Schwarz, Christian Trachsel, Martina Zanella, Claudia Riedi, Ralph Schlapbach, Alaa Othman, Can Türker, Paolo Nanni, Christian Panse

## Abstract

Mass spectrometry laboratories must turn lists of submitted samples into acquisition queues. The run order and the placement of quality-control (QC) injections determine whether a design controls batch effects and signal drift, and whether those effects stay correctable afterward. Yet operators usually set them by hand in vendor worklist editors that neither randomize run order nor offer configurable, pattern-driven QC. We present *qg*, an open-source tool that builds acquisition queues with systematic run-order handling: four run-order modes (none, simple, blocked/randomized-complete-block, and group-uniform blocked), pattern-driven QC and standard injections, and sampler- and plate-aware positioning. Unlike plate-design tools that stop at a generic sample sheet, *qg* writes the native vendor acquisition file directly, for three instrument ecosystems (Thermo Fisher XCalibur, Axel Semrau Chronos, Bruker HyStar) across proteomics, metabolomics, and lipidomics. It separates a small, stateless generation pipeline from a declarative configuration layer, so a laboratory adapts instruments, QC patterns, layouts, and naming by editing version-controlled configuration through a validating editor rather than changing code. *qg* runs from a reactive web interface or a scripted command-line interface, integrated with a LIMS (B-Fabric) or standalone from uploaded tables; randomized runs record their seed and reproduce from exported parameters. On an unbalanced design, group-uniform blocked randomization spreads biological groups evenly across acquisition time, whereas textbook block randomization leaves a tail of the largest group and can track acquisition time worse than a plain shuffle. *qg* is released under the Apache-2.0 license.

## 1 Introduction

Laboratories that run mass spectrometry (MS), whether an individual research group operating its own instrument or a laboratory serving many projects, routinely handle a steady flow of diverse sample submissions, all destined for a shared set of instruments. A core facility feels this most acutely, but the same run-order and QC decisions matter to any laboratory acquiring more than a handful of samples. Before acquiring any data, they must organize each submission into an instrument-specific queue, or worklist. This worklist is just a table, but it does a lot. It tells the system where each sample sits, which method to use, where to save the data, and how to name the files. It also fits the quality-control (QC) injections in at the right intervals. This step can seem like a simple matter of logistics, yet it has real consequences for the quality and interpretability of the final results. The order in which the instrument acquires samples, together with the frequency and type of QC injections, directly governs whether downstream analysis can detect and account for batch effects and signal drift (Cuklina et al. 2021; Dunn et al. 2011). A single run can represent weeks of instrument time and irreplaceable sample material. If a biological factor is confounded with acquisition time, no amount of post-hoc correction can fully disentangle it from instrument drift, and the whole run may yield compromised conclusions.

Two complementary safeguards address these risks. The first randomizes the run order, which prevents biological groups from aligning with acquisition time. However, simple randomization can still let a group cluster at the beginning or end by chance. Block randomization is the established remedy, though still not widespread in proteomics practice; it spreads conditions evenly across blocks, so no group is consistently favored or disadvantaged in timing (Burger et al. 2021).

The second safeguard injects pooled-QC and system-suitability samples at regular intervals (Neely et al. 2024). These injections provide repeated measurements that monitor and respond to drift. Standardized system-suitability standards further underpin longitudinal, inter-laboratory performance monitoring and harmonization across proteomics core facilities (Chiva et al. 2025). Community guidelines call for pre-planned QC and system-suitability sampling as part of study design (Broadhurst et al. 2018; Gonzalez-Dominguez et al. 2024); practical primers (Neely et al. 2024) and QC-reporting standards (Kirwan et al. 2022) support consistent implementation and reporting.

In practice, however, operators still assemble queues by hand in the worklist editors shipped with acquisition software, such as Thermo Fisher Scientific XCalibur Sequence Setup, Bruker HyStar, and Axel Semrau Chronos. These editors do not randomize run order, and their QC support, where present, is limited to fixed bracketing or calibration templating (as in XCalibur’s sequence templates) rather than the configurable, pattern-driven QC a multi-project queue needs. They are error-prone for large or multi-project batches, and they tie a laboratory to a single vendor’s table format and naming scheme. Manual editing also leaves no audit trail. The rationale for a particular run order or QC frequency lives only in the operator’s head.

A family of dedicated experimental design tools addresses the adjacent sample placement problem. OSAT allocates samples to batches so that it distributes biological groups and confounders evenly in genomics experiments (Yan et al. 2012). Omixer performs multivariate, reproducible randomization that proactively decorrelates technical and biological factors and exports lab-friendly sample sheets (Sinke et al. 2021). Well Plate Maker places samples on plates under spatial constraints using a backtracking randomized block design (Borges et al. 2021). PlateDesigner randomizes samples across microplates for multiplex immunoassays (Suprun and Suarez-Farinas 2018). PlateEditor manages complex multi-well plate layouts and associated data (Delorme et al. 2021). PLAID frames plate layout as a constraint-satisfaction problem solved by artificial intelligence (Francisco Rodriguez et al. 2023). The closest analog, InjectionDesign, provides web-based plate design with optimized stratified block randomization and QC interpolation for GC/LC–MS sample preparation. Like `qg`, it explicitly separates the physical plate layout from the injection sequence (Lu et al. 2023). Valuable as these tools are, they stop at a generic plate map or sample sheet. None emit the vendor acquisition file that a specific instrument’s control software actually loads. At the other end of the experiment, a rich ecosystem analyzes data after acquisition, including same-laboratory tooling for direct access to Orbitrap raw data and for rational LC–MS method optimization (Kockmann and Panse 2021; Trachsel et al. 2018). A practical gap remains between these two ends: a tool that couples principled run-order handling and systematic QC injection directly to multi-vendor acquisition-queue output.

We present `qg`, an open-source queue generator that closes this gap. To our knowledge, it is the only open tool that combines four run-order modes with configurable QC-injection patterns, physical sampler- and plate-aware positioning, and direct output to multiple vendor acquisition formats across proteomics, metabolomics, and lipidomics. One of those modes is a group-uniform blocked mode for unbalanced designs. Two design choices distinguish it. First, run order and QC injection are explicit, configurable inputs to the queue rather than manual steps. Experimental design best practices therefore enter the routine queue-building step. Second, every piece of site-specific knowledge lives in declarative configuration files. This knowledge covers instruments, methods, QC schemes, plate layouts, naming conventions, and vendor column mappings. An instrument scientist therefore adapts `qg` to a new instrument or workflow by editing reviewable, version-controlled configuration, with no software developer in the loop. The result is consistent, auditable queues that provide well-controlled input for downstream quantitative analysis using tools such as prolfqua and prolfquapp (Wolski et al. 2023, 2025). Because the queue fixes the acquisition order and file and folder names up front, including sample identifiers, QC labels, polarity, user, date, and instrument conventions, downstream tools can map every raw file back to its sample and model drift and batch effects against the recorded run order, rather than reconstructing them afterward.

The scale of these queues varies widely across a laboratory’s workload. A single queue may hold as few as one to four injections for instrument QC or sample-preparation method optimization. A typical two-group or multi-group comparison runs a few dozen samples (roughly 6–50). Large serum, plasma, metabolomics, or lipidomics cohorts run well over a hundred.

Queue structure varies as well. Queues routinely combine several *containers* acquired back-to-back, with QC injections separating one from the next. A *container* is the group of samples that a laboratory acquires as one unit (typically one experiment or study), identified per sample by its `container_id`. These boundaries matter for queue design, because run-order randomization and QC cadence must respect them.

## 2 Design and Implementation

### 2.1 Configuration-driven design

`qg` separates *what a laboratory runs* from *how a queue is built*. All site-specific knowledge lives in declarative configuration files (TOML and CSV), grouped by concern: `structure/` (QC-injection patterns and the QC sample register), `position/` (samplers, plate layouts, and reserved QC positions), `formatting/` (instruments, data-path templates, and vendor output formats), `methods/` (per-instrument acquisition methods), and `ui/` (interface defaults). `qg` reads configuration through a single validated accessor, `qg_configuration()`, and no other part of the application touches the configuration files directly. Adapting `qg` to a new instrument, QC scheme, plate type, or output format is therefore a configuration change, not a code change. A laboratory can version-control, review, and roll back that configuration like any other asset. Figure 1 summarises the domains of the configuration models.

**Figure 1.**
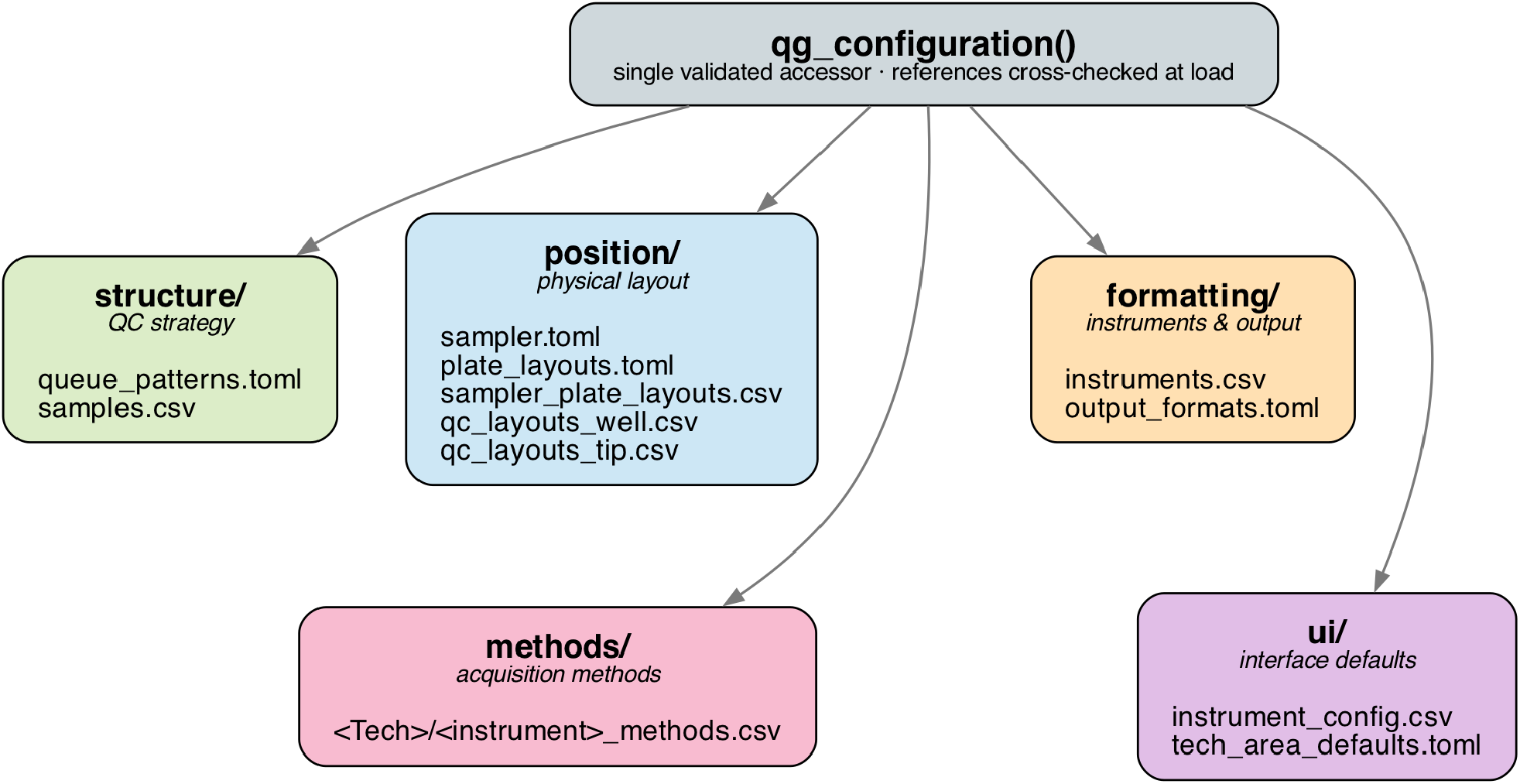
What *qg* models in configuration. A single `qg_configuration()` accessor exposes five file-based domains: QC strategy (`structure/`), physical layout (`position/`), instruments and vendor output (`formatting/`), acquisition methods (`methods/`), and interface defaults (`ui/`). It cross-checks references between these domains at load time, so a laboratory adapts *qg* by editing files, not code.

Each domain is a small declarative file. A QC-injection pattern, for example, is a TOML table in `queue_patterns.toml` listing the QC and standard injections for the `start`, `middle`, and `end` of a run and how often the middle block recurs:

~~~
[Proteomics.standard]
run_QC_after_n_samples = 8
start = [“QC03”, “QC01”]
middle = [“clean”, “QC01”]
end = [“clean”, “QC01”, “QC03”, “clean”]
separation = [“clean”, “QC01”, “clean”]
~~~

These named injections come from the per-technology sample register (`samples.csv`). Here `QC01` is a low-complexity system-suitability standard (a BSA tryptic digest with retention-time peptides, `autoQC01`), and `QC03` is a more complex performance-monitoring benchmark (K562 digest with retention-time peptides, `autoQC03`). This distinction follows the laboratory’s local autoQC nomenclature, based on the nomenclature introduced by Pichler et al. (Pichler et al. 2012); the Core for Life harmonization study provides the broader cross-laboratory context for standardized proteomics performance-monitoring materials (Chiva et al. 2025). `clean` is a buffer blank for column cleaning, and `run_QC_after_n_samples` sets how often the `middle` block recurs. The same fields carry a richer strategy for metabolomics. The `standard` metabolomics pattern brackets the run with solvent blanks (`blank`), amino-acid and organic-acid/phosphate standard mixes (`108mix_AA`, `108mix_OAP`), pooled-sample QC (`pooledQC`), and a six-step pooled-QC dilution series (`pooledQCDil1`–`pooledQCDil6`, low to high concentration). It re-injects that series every second `middle` block through the optional `middle_extended` field; the full metabolomics pattern is listed in the Supporting Information, Section S6.

Each output format is likewise a declarative column map in `output_formats.toml`, mapping internal queue fields to the vendor’s worklist columns, with literal constants and optional per-technology overlays; for example, the XCalibur SII map sets eight base columns and lets metabolomics add two more, inheriting the rest (Supporting Information, Section S6).

`qg` ships a `configuration editor`: a companion web app that exposes the same domains in a form-based interface, runs the full cross-domain validation described below on every change, and writes the underlying TOML/CSV files, either in place or as a reviewed commit. Section S2 of the Supporting Information provides a step-by-step screenshot walkthrough of the configuration editor (Figures S7–S13), covering its overview tab and each per-domain editor.

Most changes are small and local. Adjusting a QC strategy is a single edit to `queue_patterns.toml`. Changing how often QC runs is one number (`run_QC_after_n_samples`). Supporting a new plate or autosampler means defining its geometry in `plate_layouts.toml`, declaring the device in `sampler.toml`, mapping the two in `sampler_plate_layouts.csv`, and listing the reserved QC wells in `qc_layouts_well.csv`. For consumable-tip samplers such as the Evosep, list tip ranges in `qc_layouts_tip.csv` instead. Adding an instrument adds a row to `instruments.csv` and a row to the UI’s `instrument_config.csv`, together with a per-instrument acquisition-method file. Adjusting a vendor output means editing the corresponding overlay in `output_formats.toml`. `qg` also bundles an agent skill file with guided templates for these recurring tasks: standing up a new technology end-to-end, adding an instrument, adding a plate layout together with its QC samples and pattern, or adjusting a vendor output format. The skill file makes the cross-file dependencies between these tasks explicit. None of these tasks requires editing Python.

Because configuration is the contract, `qg` validates it. For example, every QC-injection pattern and every QC layout (which reserves specific plate wells or tips for QC samples) may name only samples defined in `samples.csv`, and every non-empty pattern must have at least one QC layout that provides all the QC samples it requires (Figure 2). Each file’s schema additionally enforces local invariants: injection volumes must be positive, for instance, and a file-name template may use only a fixed set of placeholders, so a mistyped placeholder is caught rather than silently accepted. `qg` thus catches these misconfigurations at load, with an explanatory message, before it generates any queue.

**Figure 2.**
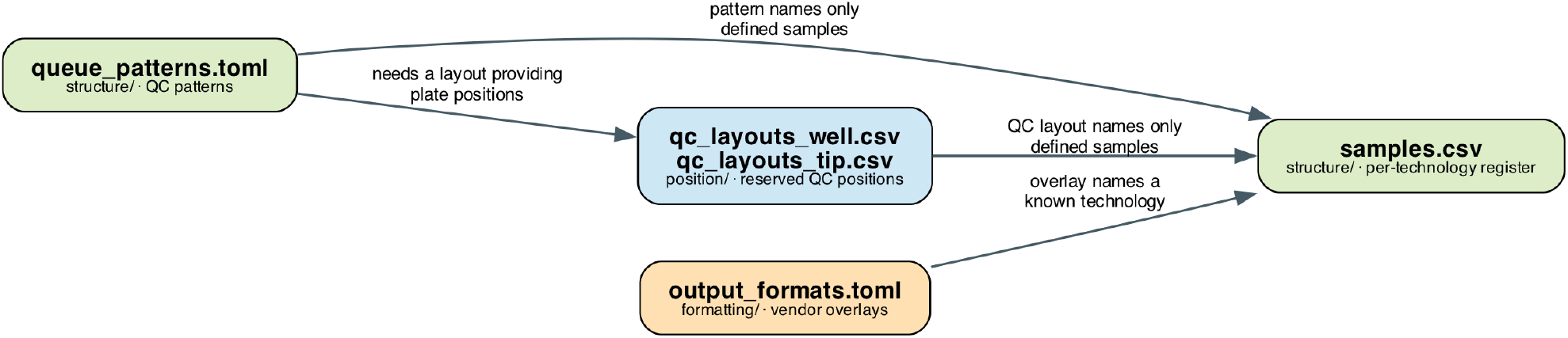
The cross-file reference constraints that `qg_configuration()` enforces at load. Each arrow points from a configuration file to the file it must be consistent with: QC patterns (`queue_patterns.toml`) and QC layouts (`qc_layouts_well.csv`/`qc_layouts_tip.csv`) may name only samples defined in `samples.csv` for the same technology; every non-empty pattern needs a QC layout that supplies the QC samples it requires; and each vendor-output overlay (`output_formats.toml`) must name a known technology. Node colours mark the configuration domain (green `structure/`, blue `position/`, orange `formatting/`). `qg` checks all of these when the configuration loads and reports every violation together, before any queue is generated.

File and folder names are themselves configuration: each instrument defines a data-path template and each sample a file-name template, both built from a fixed placeholder set (Supporting Information, Section S6). Capturing naming conventions once keeps generated names consistent, so downstream QC and reporting tools can trace every raw file to its LIMS sample.

### 2.2 The generation pipeline

Queue generation is a short sequence of pure functions that transform the structured inputs into a vendor table (Figure 3). The two inputs are the queue parameters and the sample list. The queue parameters fix everything a run needs beyond the samples themselves. They name the technology, instrument, sampler, and plate and QC layouts. They set the QC pattern, the output format, and the per-polarity acquisition methods. They give the randomization mode and, optionally, its seed. They also supply the user and date that build the data paths. The sample list provides, for each sample, a name, a stable identifier, the container it belongs to (its `container_id`), and an optional grouping variable for blocked randomization. For plate submissions, each sample also includes its plate and well position. Vial and plate submissions are both converted to the same positioned format before randomization proceeds. The paragraphs below describe the stages in the same order as Figure 3.

**Figure 3.**
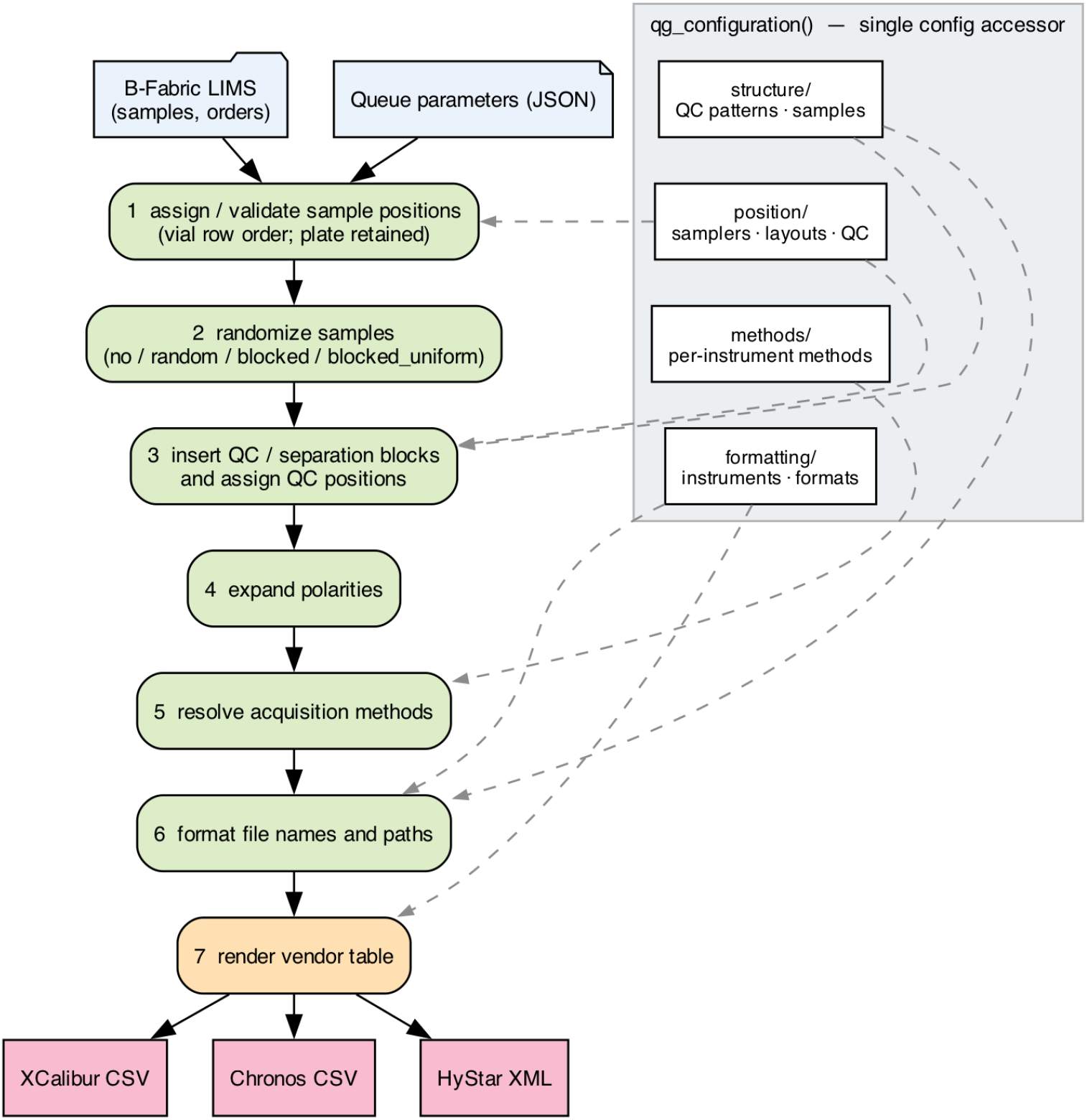
Two-tier architecture of *qg*: a stateless seven-step generation pipeline (centre) over a declarative, config-driven foundation read through a single `qg_configuration()` accessor (right). Sample metadata and queue parameters enter at the top. The pipeline assigns or validates sample positions, randomizes user samples, inserts QC and separation blocks with their reserved QC positions, expands requested polarities, resolves acquisition methods, formats file names and paths, and renders the vendor table. Dashed arrows show which configuration group feeds each stage.

#### User-sample positioning

User samples enter `qg` either as a plate queue or as a vial queue. In plate mode, the sample list already contains plate and well positions, which `qg` validates against the sampler layout and the reserved QC positions and then retains. In vial mode, samples arrive without physical positions. `qg` assigns those positions at the start of generation, before any run-order randomization, by filling the available sampler positions in row-major order from the selected start tray and start position. The resulting positioned queue is the object that enters randomization and all later generation stages.

The current row-order allocation is intentional. A group-aware placement optimizer could distribute biological groups across the plate, but without robotic support it would create more complex manual transfer schemes and a higher risk of loading errors. Such plate allocation is therefore better treated as a sample-preparation or automation step, while `qg` keeps manual vial-to-plate loading simple and reproducible.

#### Randomization

`qg` supports four modes. `no` keeps submission order, which suits samples that were already randomized upstream or whose ordering is externally fixed. `random` shuffles samples within each plate and container. `blocked` implements a randomized complete-block design (RCBD). It groups samples by a user-specified grouping variable (typically the biological condition), shuffles within each group, and emits one member of each group per block, so the run interleaves the conditions rather than clustering them in time (Burger et al. 2021). When the groups are *unbalanced*, RCBD exhausts the smaller groups first and leaves a tail dominated by the largest group. `blocked_uniform` removes this artifact: a fair-share rule spreads each group as evenly as possible across the whole run, so even a dominant group does not concentrate at the end. InjectionDesign applies a related group-uniform interpolation for unbalanced designs (Lu et al. 2023); `qg` makes it an explicit, named run-order mode wired directly to vendor output. Crucially, all four modes shuffle only *within* plate and container boundaries. The randomization therefore never violates physical layout constraints or the separation between different containers. In a multi-container queue, randomization stays within each container. A container’s samples remain contiguous rather than interleaved across the whole run. The instrument therefore acquires them close together in time, which confines drift within a container rather than spreading it across the entire queue.

#### QC-injection patterns

During generation, `qg` turns the selected declarative pattern into concrete queue rows. It inserts the start block before the first container, separation blocks between containers, middle blocks within each container after the configured number of user samples, optional extended blocks at their configured recurrence, and the end block after the last container. This is the pipeline-stage use of the `queue_patterns.toml` schema described above in the configuration-driven design section, not a second definition of that schema. Periodic pooled-QC at a set interval follows established metabolomics practice (Dunn et al. 2011; Broadhurst et al. 2018; Kirwan et al. 2022); re-injecting a dilution series throughout the run extends Broadhurst’s one-off response-linearity check (Broadhurst et al. 2018) into repeated monitoring, a `qg` design choice. The lipidomics `standard` pattern brackets the run with an EquiSPLASH class-spanning internal-standard mix (one labeled standard per lipid class, a principle validated across nine LC–MS platforms (Cajka et al. 2017)) together with a pooled-QC dilution series, and acquires every sample in both ionization polarities, in line with community lipidomics QC practice (Burla et al. 2018); a worked lipidomics queue is shown in the Supporting Information, Figure S16. Because the pattern is data, a laboratory can encode several strategies side by side and select between them per queue. One scheme can be dense for method development, and another sparse for routine runs.

#### QC positioning

After the QC and separation blocks have been inserted, `qg` assigns QC and standard injections to physical positions defined by the selected QC layout. Well-plate samplers, such as the Thermo Vanquish and Waters M-Class, and tip-based samplers, such as the Evosep, use different position providers, but the principle is the same: the QC layout reserves named wells or tips for the QC samples declared by the selected pattern. Recurring QC injections therefore return to the same physical positions regardless of the order of user samples (Figure 4). This separates physical layout from injection sequence: the operator loads the plate once, in a stable arrangement, while `qg` generates the acquisition order independently.

**Figure 4.**
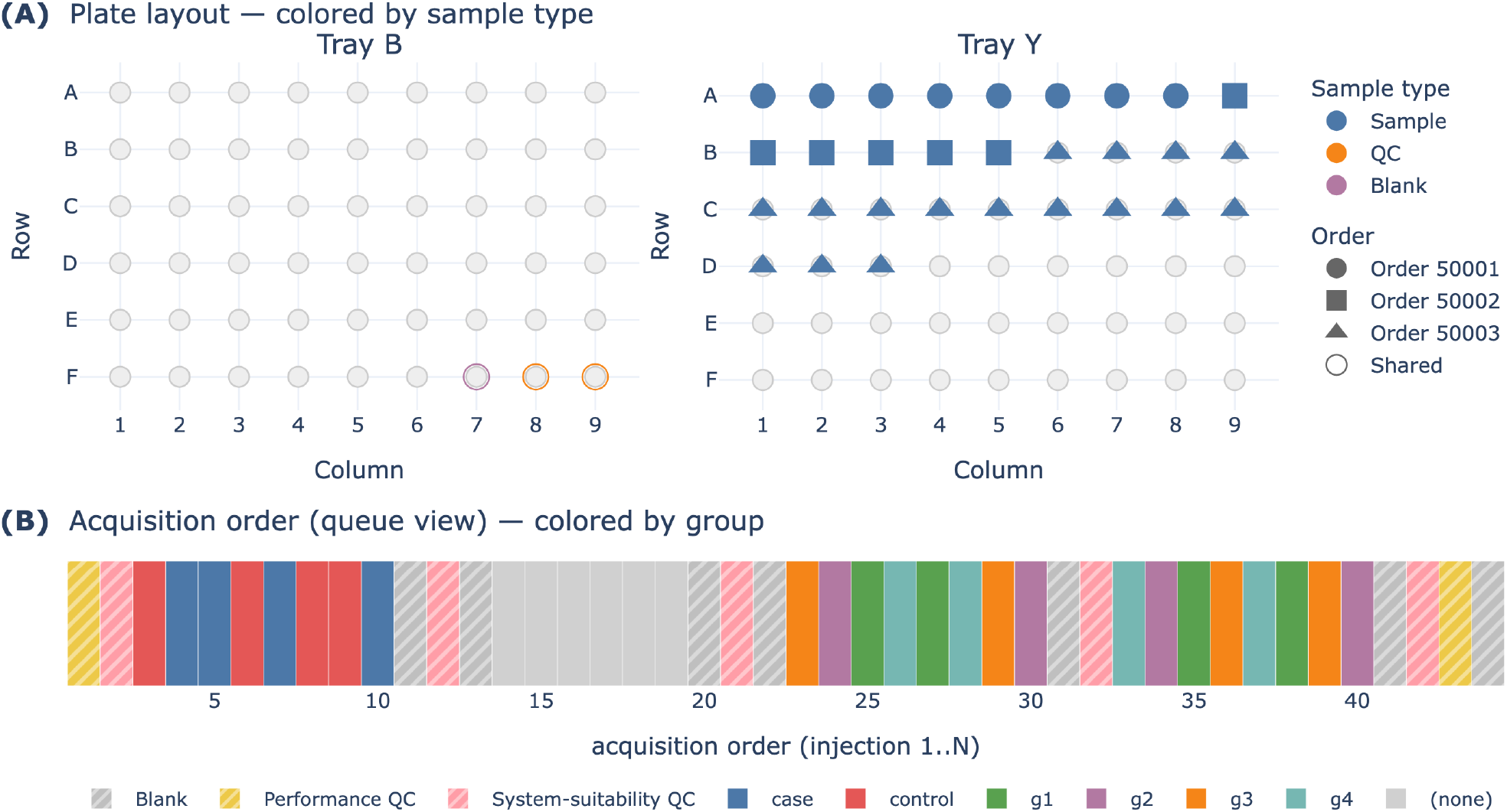
A representative multi-container proteomics queue (three orders of 8, 6, and 16 samples, for 30 user samples and 44 injections in total; Thermo Vanquish, `standard` QC pattern), rendered by the same routines the web interface uses. `(A)` Plate layout colored by sample type, with marker shape encoding the order: user samples fill the sample tray, while QC and cleaning injections occupy reserved QC positions on a dedicated tray, including the `separation` blocks inserted between consecutive orders. `(B)` The same queue as an acquisition-order timeline (one tile per injection along the run) colored by biological group. The run uses `blocked_uniform` randomization, so within each order the timeline interleaves the groups across its injections rather than acquiring them in contiguous blocks. QC and blank injections are grey, and the one order that carries no grouping variable appears as “(none)”. Both panels render from the very same queue object that *qg* writes to disk.

#### Polarity, methods, and naming

Once the positioned injection sequence is fixed, `qg` adds acquisition-specific detail. If a run requires more than one ionization polarity, the generator duplicates each injection for the requested polarities and assigns the final run numbers after that expansion. It then resolves the acquisition method for every user-sample and QC injection from the selected technology, instrument, sample type, polarity, and method map. File names and data paths are rendered from templates using recorded queue fields, including run number, container identifier, sample identifier, sample name, polarity, method name, date, and user. When configured, `qg` also marks the last injection of each container subqueue in the file name, so downstream QC can recognize container boundaries.

#### Vendor output

A final mapping step renders the rows into the requested format: Thermo XCalibur CSV, Axel Semrau Chronos CSV, or Bruker HyStar XML. Each vendor format is declared as a column map in `output_formats.toml`; the same mechanism also supports technology-area additions, with the full configuration example shown in the Supporting Information, Section S6. The renderer applies format-specific position conversion, such as flattening alphanumeric wells to Evosep vial indices, and adding a supported format is configuration-only when it can reuse an existing serializer; genuinely new wire formats, such as Bruker HyStar XML, require a writer function.

### 2.3 Interfaces and deployment

The operator uses `qg` through a reactive web interface built with marimo (Figure 5). Cascading menus let the operator select technology, instrument, sampler, plate and QC layouts, pattern, and randomization mode, and they only ever offer valid combinations. An editable table shows the incoming sample list. There the operator can deselect samples from the run and correct their grouping variable and other metadata.

**Figure 5.**
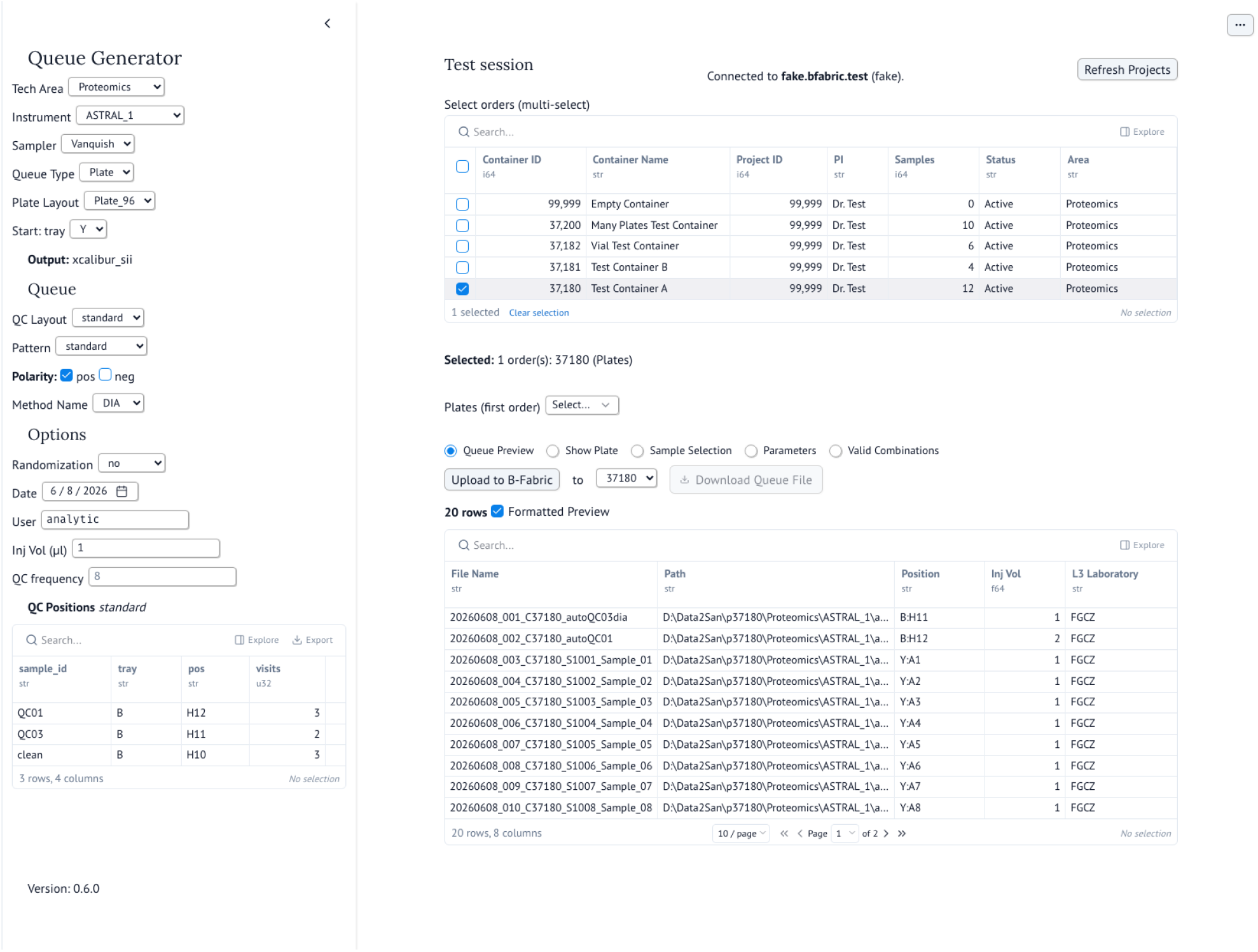
The *qg* web interface. Cascading menus (left) constrain the operator to valid instrument/sampler/layout/pattern combinations; the sample list is editable and reorderable; and the formatted queue and plate map preview update reactively. The previewed queue is the exact table written to disk.

When randomization is off, or when an externally fixed order must be preserved, the operator can also reorder samples manually. The interface previews both the formatted queue and a plate map. The operator then downloads the queue file, or the run parameters as a single JSON document. A complete queue-app walkthrough is provided in the Supporting Information, Section S1 (Figures S1–S6). Because the menus draw on the same validated configuration that drives generation, a well-formed configuration cannot lead the operator to an invalid run.

`qg` also exposes the same generation core as a command-line interface that consumes a JSON file of parameters. This enables scripted, batch, and continuously tested runs without the GUI. In production, the application integrates with the B-Fabric LIMS (Panse et al. 2022), which links projects, orders, samples, instruments, applications, storage, and work units into one audited data-management layer. `qg` reads sample metadata for a selected order and deposits the generated queue back as a work unit, so the queue stays linked to its samples for later audit. A single code path serves two access levels: facility staff can browse all orders, while external users see only their own. Both levels share an identical generation core. For groups without a LIMS, the same application also runs standalone. The operator uploads a CSV or XLSX sample table, configures the queue with identical controls, and downloads the result locally. The uploaded table uses the same normalized schema that `qg` derives from a B-Fabric order (Supporting Information, Section S3, Table S1), so a hand-prepared spreadsheet and a LIMS export build a queue identically. `qg` deploys the web application and the configuration editor from a Git repository, so a laboratory reviews each configuration change as a commit, and the change reaches every operator at once.

## 3 Results and Discussion

### 3.1 A representative queue

Figure 4 shows a queue generated for a representative proteomics run. The run combines three orders acquired back to back, of 8, 6, and 16 samples, 30 in total, on a Thermo Vanquish well plate using the `standard` QC pattern. The engine produced 44 injections: the 30 user samples plus 14 QC injections. Those QC injections come from the pattern’s start, middle, and end blocks and from the `separation` blocks inserted between consecutive orders. Panel (A) shows the resulting plate, colored by sample type: the user samples fill the sample tray, while QC and cleaning injections occupy the reserved QC positions on a dedicated tray. Panel (B) shows the same queue as an acquisition-order timeline colored by biological group. The run uses `blocked_uniform` randomization, so within each order the timeline interleaves the groups across its injections rather than acquiring them in contiguous blocks. One order carries no grouping variable and appears as “(none)”. Each order’s samples remain contiguous and QC-separated, so the multi-container queue respects container boundaries while sharing a single physical layout. The two panels therefore show the generated run itself, once as a physical layout and once as an acquisition-order timeline.

### 3.2 Run-order randomization on an unbalanced design

Randomization aims to remove the association between a sample’s biological group and its position in the acquisition order, so that instrument drift does not masquerade as a biological effect. We generated a queue of 56 samples drawn from four *unbalanced* biological groups (24, 14, 10, and 8 samples) and applied each randomization mode under a fixed seed (Figure 6). This comparison shows what `qg`’s modes achieve, and why an unbalanced design needs more than textbook RCBD. To quantify the residual confounding, we computed *η*^2^, the share of variance in acquisition position that biological group explains (the between-group sum of squares divided by the total, as in one-way ANOVA). *η*^2^ runs from 0, when group and acquisition order are unrelated, to 1, when the group is perfectly aliased with time.

**Figure 6.**
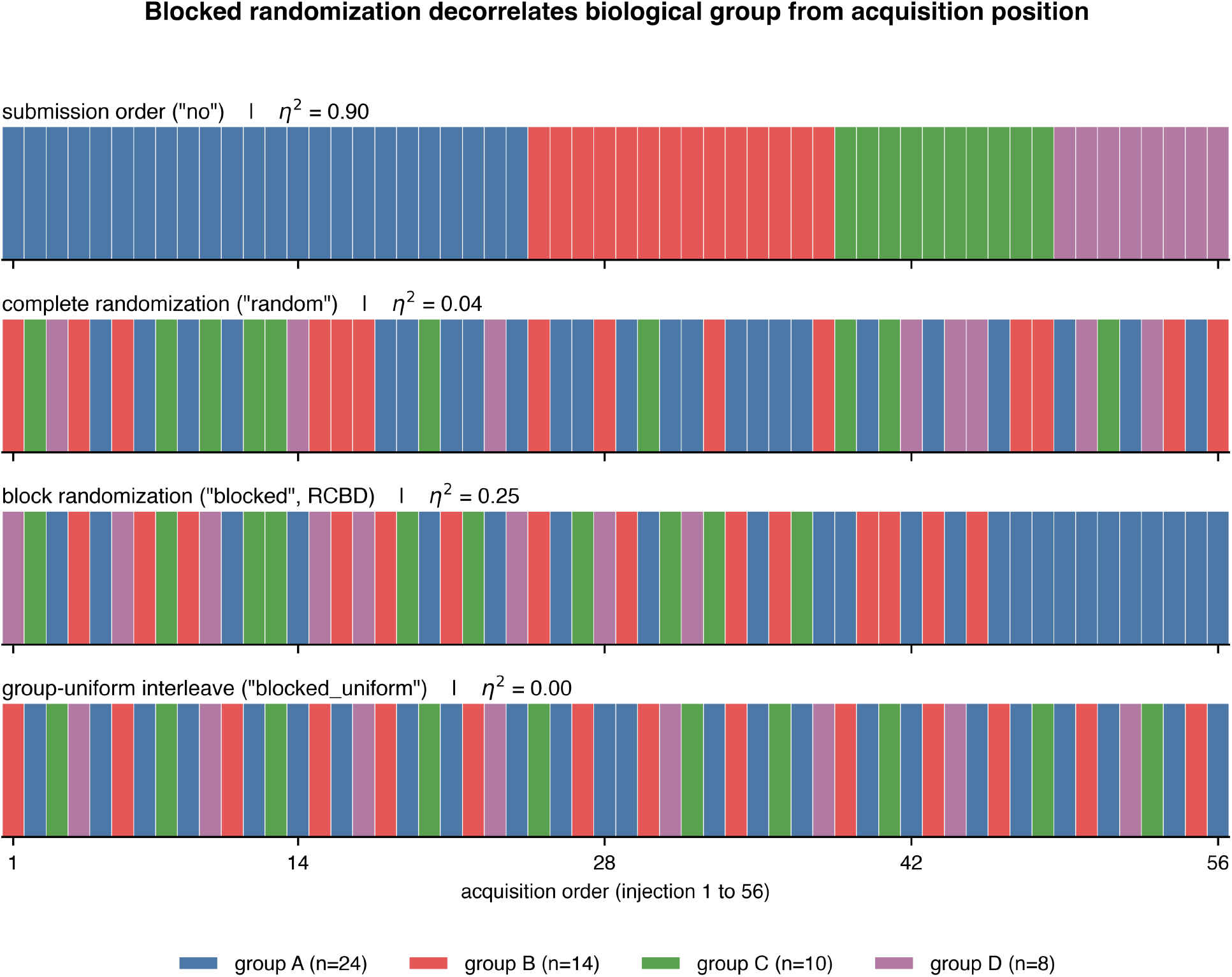
Randomization removes group–time confounding, and for unbalanced groups the mode matters. For 56 samples in four unbalanced groups (24, 14, 10, 8; colours), each panel shows the acquisition order (left to right) under one mode. Submission order (`no`) places groups in contiguous blocks; complete randomization (`random`) interleaves them but leaves local clustering; RCBD block randomization (`blocked`) places one sample of each group per block but, once the smaller groups are spent, leaves a tail of the largest group; group-uniform interleaving (`blocked_uniform`) spreads every group evenly across the whole run. The per-panel statistic is *η*^2^, the share of acquisition-position variance explained by group (lower is better): 0.90, 0.04, 0.25, and 0.00 respectively.

With no randomization, the four groups occupy contiguous blocks, and the association is, by construction, near-maximal: biological group explains 90% of the variance in acquisition position (*η*^2^ = 0.90), so the group is essentially aliased with time. Complete randomization cuts this to *η*^2^ = 0.04, but, as expected for an unconstrained shuffle, it leaves visible local clustering. The RCBD mechanism described above improves interleaving for balanced groups but not for this unbalanced queue: `blocked` gives *η*^2^ = 0.25, worse than a plain shuffle, because its majority-group tail shifts that group’s mean position late. `blocked_uniform` instead drives *η*^2^ to essentially zero (0.002), far below RCBD. For unbalanced designs, the ordering is therefore `blocked_uniform` ≪ `random` < `blocked` ≪ `no`. These rankings are stable over 2000 seeds, and a second metric, the longest consecutive run of one group, gives the same ordering (≈2 for `blocked_uniform`, ≈4 for `random`, ≈11 for RCBD); an imbalance sweep confirms that RCBD overtakes a plain shuffle once the dominant group exceeds roughly twice the others (Supporting Information, Section S7, Figures S14 and S15).

### 3.3 Comparison with existing tools

Table 1 positions `qg` against the sample-placement and plate-design tools discussed above. It extends the feature comparison of Lu et al. (Lu et al. 2023) with a `qg` column and with the acquisition-specific features that distinguish it. The plate-design axes cover the web service, the graphical interface, addressing batch effects, QC sample support, and the separation of physical layout from injection sequence. On these axes, `qg` matches the capabilities of the most capable existing tools, including InjectionDesign; InjectionDesign even offers a richer interactive plate-layout designer than `qg`. The decisive difference is on the acquisition-queue axes. `qg` is the only tool in the set that writes a native MS acquisition file^1^, and it does so for three vendor ecosystems from one internal queue. The others stop at a generic plate map or sample sheet that an operator must still transcribe into the instrument’s worklist editor. `qg` additionally expands acquisition polarity for metabolomics and lipidomics, integrates with a LIMS, and runs from the same deterministic core through both a GUI and a CLI. The QC model also differs from the closest analog. InjectionDesign spaces QC injections automatically by inserting them evenly in a circular fashion, whereas `qg` places them according to an explicit, version-controlled pattern, so a laboratory states exactly which QC injections appear and where. Because `qg` also runs without a LIMS, it needs neither an upstream plate-design tool nor B-Fabric to produce a vendor queue. The comparison is therefore complementary rather than adversarial. A laboratory could reasonably use a plate-design tool to lay out a plate and `qg` to build the instrument queue. For the queue-building step specifically, though, `qg` covers ground the others leave to manual editing.

**Table 1.** Feature comparison of *qg* with sample-placement and plate-design tools. The upper block adapts the feature axes of Lu et al. (their Table 1), extended with a *qg* column and corrected against each tool’s publication; the lower block adds acquisition-queue features assessed from each tool’s publication. *QC sample support* here means a tool can place control or QC samples within the layout, not that it schedules QC injections through the run. ✓ = yes, ✗ = no, ? = not documented.

|  | WPM | PlateEditor | PlateDesigner | PLAID | Omixer | OSAT | InjectionDesign | *qg* |
| --- | --- | --- | --- | --- | --- | --- | --- | --- |
| *Plate / sample-prep features (after Lu et al.)* |  |  |  |  |  |  |  |  |
| Web service | ✗ | ✓ | ✓ | ✓ | ✗ | ✗ | ✓ | ✓ |
| Graphical interface | ✓ | ✓ | ✓ | ✓ | ✗ | ✗ | ✓ | ✓ |
| Addresses batch effects | ✓ | ✗ | ✓ | ✓ | ✓ | ✓ | ✓ | ✓ |
| QC sample support | ✓ | ✓ | ✓ | ✓ | ✓ | ✗ | ✓ | ✓ |
| Layout/sequence separation | ✗ | ✗ | ✗ | ✗ | ✗ | ✗ | ✓ | ✓ |
| Customizable worksheet | ✗ | ✗ | ✗ | ✗ | ✗ | ✗ | ✓ | ✓ |
| *Acquisition-queue features* |  |  |  |  |  |  |  |  |
| Native MS acquisition file | ✗ | ✗ | ✗ | ✗ | ✗ | ✗ | ✗ | ✓ |
| Multiple vendor formats | ✗ | ✗ | ✗ | ✗ | ✗ | ✗ | ✗ | ✓ |
| CLI / scriptable | ✓ | ✗ | ✗ | ✓ | ✓ | ✓ | ✗ | ✓ |
| Polarity expansion | ✗ | ✗ | ✗ | ✗ | ✗ | ✗ | ? | ✓ |
| LIMS integration | ✗ | ✗ | ✗ | ✗ | ✗ | ✗ | ? | ✓ |

### 3.4 Coverage and extensibility

The configuration shipped with `qg` already spans a realistic core-facility instrument park: 15 instruments across three technologies (nine proteomics, three metabolomics, three lipidomics; these counts exclude an internal test-fixture technology), three autosamplers (the well-plate Vanquish and M-Class and the tip-based Evosep), three plate layouts, and four vendor output formats that resolve to three writers. The QC layer is equally concrete: eleven named QC-injection patterns and thirty QC and standard-injection definitions, besides the per-technology `default` user-sample template. These counts are tabulated in the Supporting Information, Section S5 (Tables S2 and S3). The entries range from simple proteomics cleaning-plus-QC brackets to metabolomics pooled-QC dilution series and chemical-standard mixes. None of these is hard-coded. Each is a row or table in a configuration file, and the same files are the extension points. A laboratory onboarding a new Orbitrap, a new plate geometry, or a vendor variant adds entries to the relevant files. The instrument then appears throughout without a software release: in the interface menus, in method resolution, and in the generated queue. This is the practical payoff of the configuration-driven design. The people who know the instruments and the QC requirements own the adaptation, rather than the people who maintain the code. They work through the configuration editor and bundled assisted-editing skill described above, making each adaptation reviewable and version-controlled.

### 3.5 Reproducibility

`qg` builds in reproducibility at several levels. For a given input, the non-randomized pipeline is fully deterministic. The randomized modes draw from a seeded generator, and the seed is part of the run parameters. When `qg` generates a queue, it records the seed actually used back into the exported parameters. The operator may supply that seed, or `qg` may draw it automatically when none is given. Either way, a randomized run reproduces exactly from its own record, rather than depending on a separately remembered value. The complete set of run parameters covers the technology, instrument, sampler, layouts, pattern, randomization mode, seed, and the sample list. `qg` exports this set as a single JSON document, and the command-line interface regenerates the queue from that document without the GUI; a worked JSON-and-CLI reproduction is shown in the Supporting Information, Section S4. A laboratory can therefore archive, share, and reproduce a run from one file, and `qg` attaches the same parameters to the B-Fabric work unit. Because the engine runs exactly once per queue, the interface preview, the downloaded file, a CLI run, and the deposited work unit all derive from one generated queue object. The downloaded vendor file is meant to be an output artifact, not a surface for hand editing. `qg` decides and records sample selection and order, QC scheme, and naming upstream, so manual post-editing of the worklist becomes unnecessary. Finally, an automated end-to-end test reproduces a full interface session and checks the downloaded queue byte-for-byte against a stored reference, so continuous integration flags any change that would alter the generated queues. Together, these levels make a queue a reproducible function of recorded inputs rather than a one-off manual artifact. Two further properties make the queues trustworthy in practice. First, `qg` validates each queue as it is generated: it checks that the chosen sampler and plate layout have capacity for all injections, and flags any collision between reserved QC positions and user-sample placement, so an over-capacity or mis-placed queue is rejected before output rather than discovered on the instrument. Second, `qg` is a mature production tool rather than a prototype: its lineage has been in routine use at the FGCZ since 2015 (see Author Contributions), and the configuration-driven version described here generates the facility’s acquisition queues across all three vendor ecosystems, with its output validated on production instruments during normal operation.

## 4 Limitations and Future Work

`qg` deliberately scopes itself to the queue-building step, and several boundaries follow from that. Its randomization balances and shuffles within plate and container boundaries. It does not optimize the allocation of samples across plates or batches to balance covariates. The block designs balance a single grouping variable. `qg` does not yet support multi-factor stratification, which some randomization tools offer. Output currently covers three vendor ecosystems. Broadening the set of acquisition formats and adding samplers are ongoing efforts, and they are largely configuration and isolated extension rather than structural changes. These boundaries are intentional. `qg` also schedules and writes QC and standard injections but does not interpret them: evaluating QC results and accepting or rejecting a run are downstream steps, outside its scope. Within the queue-building step itself, the behavior is complete and recorded: run order, QC placement, physical positioning, naming, and reproducibility.

## 5 Conclusions

`qg` turns plain sample lists into vendor-ready acquisition queues for the acquisition step that lies between upstream experimental-design tools and downstream QC monitoring. It writes the actual instrument worklist and records the decisions that otherwise tend to remain manual: run order, QC schedule, physical positions, naming, and output format. Site-specific instruments, QC schemes, and naming conventions stay in version-controlled configuration that an instrument scientist can maintain without code changes. Blocking acquisition order to control group–time confounding is established design practice; our contribution is to make it executable in the vendor queue, including a group-uniform fair-share mode for unbalanced designs where textbook block randomization can underperform (Zhao and Berger 2017). By making run order and QC injection reproducible and auditable properties of the queue, `qg` provides well-controlled input to downstream quantitative analysis. In practice this means fewer manual transcription errors, less operator time spent assembling worklists, and consistent run-order and QC design at routine queue construction.

## 6 Data and Code Availability

`qg` is open-source under the Apache-2.0 license. The source code is available from its public repository (https://github.com/fgcz/qg), and each release is archived with citation metadata on Zenodo under a persistent DOI (Wolski et al. 2026). A public browser-based demo instance, which requires no installation or account, is available at https://apps-dev.bfabric.org/queue-gen-local/. The computational provenance for the manuscript and SI figures is summarized in Section S9 of the Supporting Information.

## 7 Acknowledgements

The authors thank Maria Di Errico, Jonas Grossmann, Sibylle Pfammatter, Annika Jagels, Tobias Kockmann, and Bernd Roschitzki for valuable discussions and feedback.

## 8 Author Contributions

Conceptualization: Christian Trachsel, Christian Panse, Witold E. Wolski, Leonardo Schwarz. Requirements specification: Paolo Nanni, Alaa Othman, Martina Zanella, Claudia Riedi. LIMS integration model, B-Fabric sample-input workflow, and workunit deposition: Can Türker, Christian Panse, Leonardo Schwarz. Software: Christian Trachsel (first implementation, 2015), Christian Panse (a decade of maintenance and extension), Witold E. Wolski and Leonardo Schwarz (2026 AI-assisted refactor). Software architecture, configuration system, randomization strategy, and GUI design: Witold E. Wolski, Leonardo Schwarz. Validation and user testing: Martina Zanella, Claudia Riedi. Supervision, resources, and funding acquisition: Ralph Schlapbach. Writing – original draft: Witold E. Wolski. Writing – review and editing: all authors.

## 9 AI Use Disclosure

We used AI-based tools extensively between December 2025 and July 2026, in an agentic, AI-assisted development and writing workflow. Several large language models assisted with software refactoring, configuration design, test generation, and the drafting and editing of this manuscript: Anthropic Claude (Opus 4.5–4.8 and Sonnet 4.6), OpenAI Codex (5.4–5.5), Mistral Large 3, and Devstral 2. We used Grammarly to review and correct the text. These tools do not meet authorship criteria, and we do not list them as authors. We directed all AI use and reviewed the resulting code, tests, and text; in particular, the AI-generated test suite and the configuration-validation logic that underpin the tool’s reliability were reviewed and verified by the authors. We take full responsibility for the content of this article. The complete source code and configuration are openly available (see Data and Code Availability).

## Footnotes

1 PlateDesigner and PLAID do emit native files for plate readers and liquid-handlers (xPONENT; ECHO, I.DOT), but not MS acquisition files.

