## Supplemental Data 1 for "*qg*: Configuration-Driven, Multi-Vendor Acquisition Queue Generation with Reproducible Run-Order and QC Control for Mass Spectrometry"

#### 1 Queue-app walkthrough

This section follows one operator session in the `qg` web interface, from order selection to a downloadable queue, for the example order shown throughout. The cascading parameter menus, the editable sample table, the formatted queue preview, and the plate/timeline visualizations are all driven by the same validated configuration and the same generation core described in the main text.

The operator first selects an order and its samples (Figure 1), then sets the run parameters through cascading menus that only ever offer valid instrument/sampler/layout/pattern combinations (Figure 2).

The sample table is editable (Figure 3): samples can be deselected from the run, their grouping variable and other metadata corrected, and — when randomization is off or an externally fixed order must be preserved — reordered by hand. The formatted queue and a plate map update reactively, and the preview is byte-for-byte the file that downloads (Figure 4).

Two visualizations help the operator judge the run before acquisition. The plate map (Figure 5) colours wells by group and reports a group $\leftrightarrow$ plate-position balance score; the acquisition timeline (Figure 6) shows the injection sequence with its group $\leftrightarrow$ run-position balance.

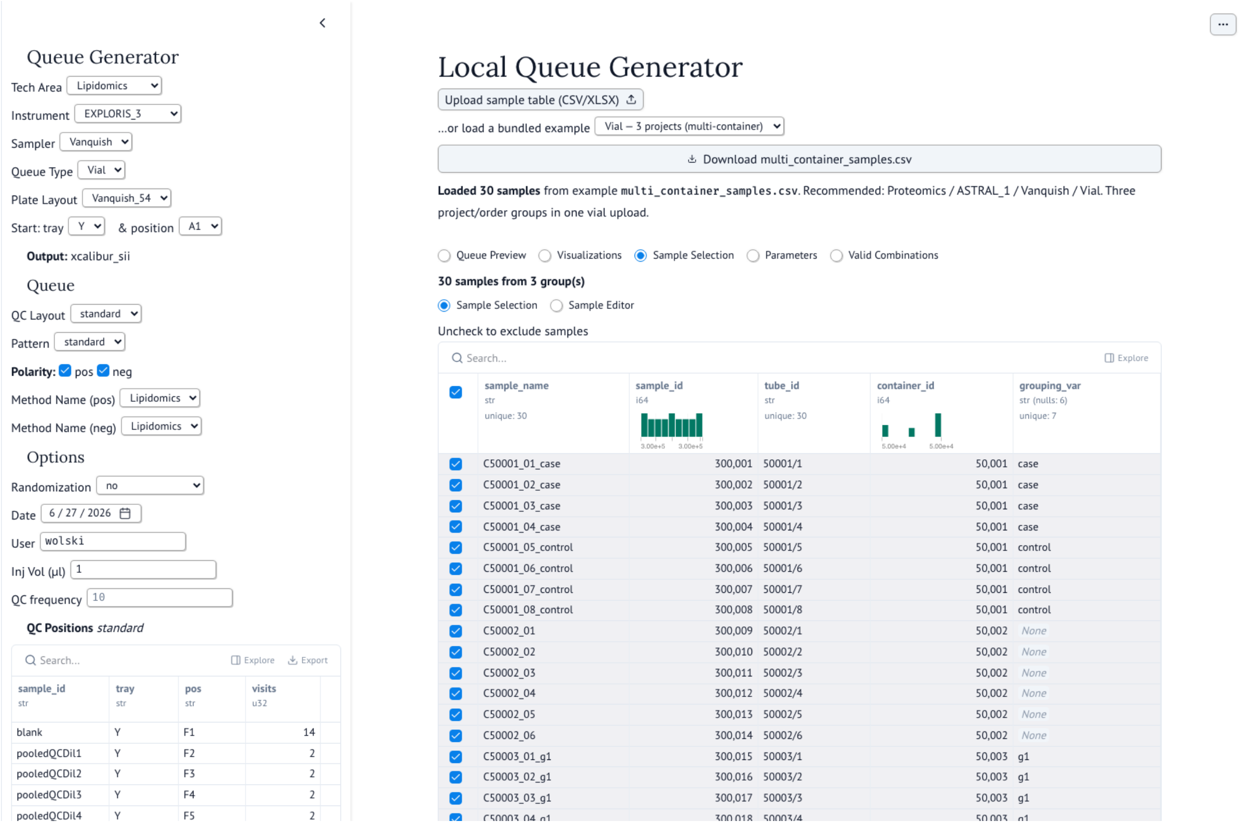


Figure 1: Order and sample selection. The operator picks one or more projects/containers; the incoming samples are listed for review before configuration.

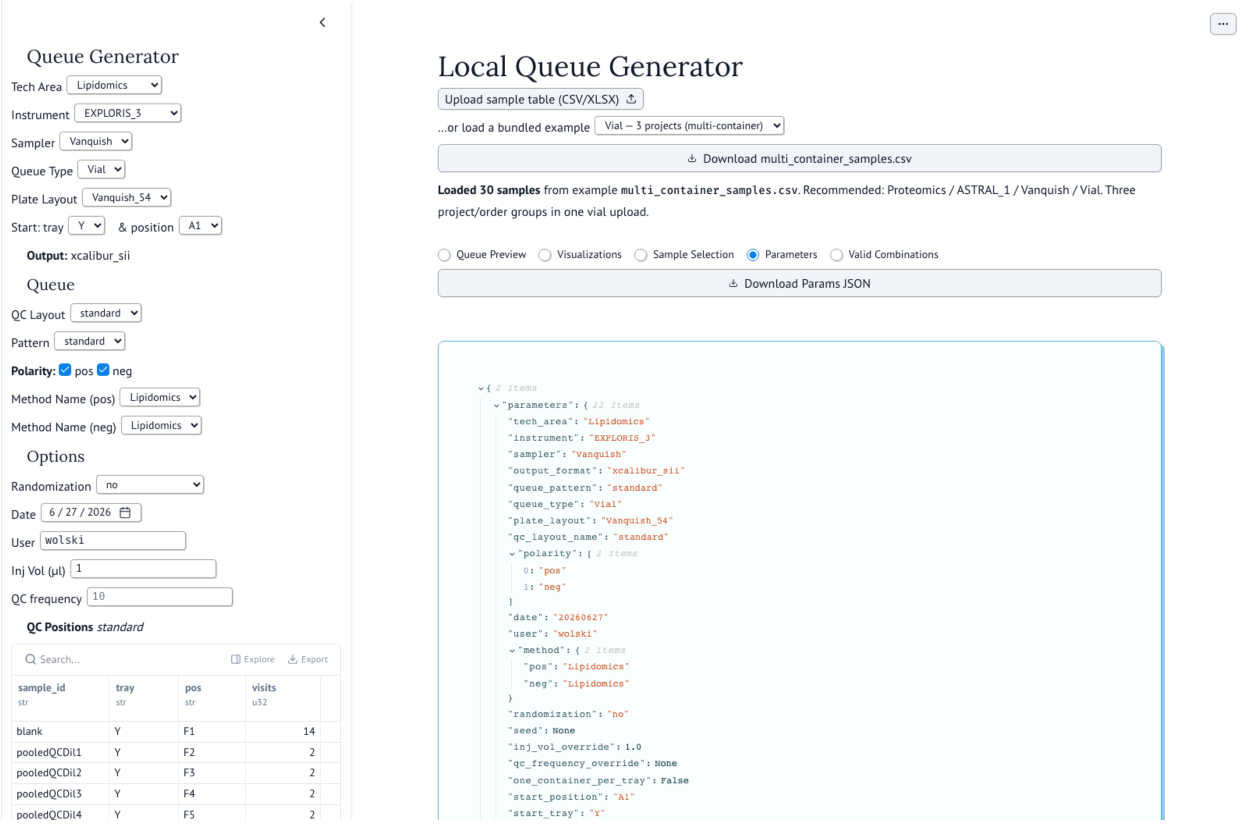


| sample_id | tray | pos | visits |
|---|---|---|---|
| blank | Y | F1 | 14 |
| pooledQCDil1 | Y | F2 | 2 |
| pooledQCDil2 | Y | F3 | 2 |
| pooledQCDil3 | Y | F4 | 2 |
| pooledQCDil4 | Y | F5 | 2 |


```
{ 2 items
  "parameters": { 22 items
    "tech_area": "Lipidomics"
    "instrument": "EXPLORIS_3"
    "sampler": "Vanquish"
    "output_format": "xcalibur_sii"
    "queue_pattern": "standard"
    "queue_type": "Vial"
    "plate_layout": "Vanquish_54"
    "qc_layout_name": "standard"
    "polarity": [ 2 items
      0: "pos"
      1: "neg"
    ]
    "date": "20260627"
    "user": "wolski"
    "method": { 2 items
      "pos": "Lipidomics"
      "neg": "Lipidomics"
    }
    "randomization": "no"
    "seed": None
    "inj_vol_override": 1.0
    "qc_frequency_override": None
    "one_container_per_tray": False
    "start_position": "A1"
    "start_tray": "Y"
```


Figure 2: Run parameters. Cascading menus expose only valid combinations of technology, instrument, sampler, plate and QC layouts, QC pattern, randomization mode, and output format; these are exactly the queue parameters serialised to the run's JSON.

| ☑ | sample_name<br>str<br>unique: 30 | sample_id<br>i64<br>3.00e+5 3.00e+5 | tube_id<br>str<br>unique: 30 | container_id<br>i64<br>5.00e+4 5.00e+4 | grouping_var<br>str (nulls: 6)<br>unique: 7 |
|---|---|---|---|---|---|
| ☑ | C50001_01_case | 300,001 | 50001/1 | 50,001 | case |
| ☑ | C50001_02_case | 300,002 | 50001/2 | 50,001 | case |
| ☑ | C50001_03_case | 300,003 | 50001/3 | 50,001 | case |
| ☑ | C50001_04_case | 300,004 | 50001/4 | 50,001 | case |
| ☑ | C50001_05_control | 300,005 | 50001/5 | 50,001 | control |
| ☑ | C50001_06_control | 300,006 | 50001/6 | 50,001 | control |
| ☑ | C50001_07_control | 300,007 | 50001/7 | 50,001 | control |
| ☑ | C50001_08_control | 300,008 | 50001/8 | 50,001 | control |
| ☑ | C50002_01 | 300,009 | 50002/1 | 50,002 | None |
| ☑ | C50002_02 | 300,010 | 50002/2 | 50,002 | None |
| ☑ | C50002_03 | 300,011 | 50002/3 | 50,002 | None |
| ☑ | C50002_04 | 300,012 | 50002/4 | 50,002 | None |
| ☑ | C50002_05 | 300,013 | 50002/5 | 50,002 | None |
| ☑ | C50002_06 | 300,014 | 50002/6 | 50,002 | None |
| ☑ | C50003_01_g1 | 300,015 | 50003/1 | 50,003 | g1 |
| ☑ | C50003_02_g1 | 300,016 | 50003/2 | 50,003 | g1 |
| ☑ | C50003_03_g1 | 300,017 | 50003/3 | 50,003 | g1 |
| ☑ | C50003_04_g1 | 300,018 | 50003/4 | 50,003 | g1 |
| ☑ | C50003_05_g2 | 300,019 | 50003/5 | 50,003 | g2 |
| ☑ | C50003_06_g2 | 300,020 | 50003/6 | 50,003 | g2 |
| ☑ | C50003_07_g2 | 300,021 | 50003/7 | 50,003 | g2 |
| ☑ | C50003_08_g2 | 300,022 | 50003/8 | 50,003 | g2 |
| ☑ | C50003_09_g3 | 300,023 | 50003/9 | 50,003 | g3 |
| ☑ | C50003_10_g3 | 300,024 | 50003/10 | 50,003 | g3 |
| ☑ | C50003_11_g3 | 300,025 | 50003/11 | 50,003 | g3 |
| ☑ | C50003_12_g3 | 300,026 | 50003/12 | 50,003 | g3 |
| ☑ | C50003_13_g4 | 300,027 | 50003/13 | 50,003 | g4 |
| ☑ | C50003_14_g4 | 300,028 | 50003/14 | 50,003 | g4 |
| ☑ | C50003_15_g4 | 300,029 | 50003/15 | 50,003 | g4 |
| ☑ | C50003_16_g4 | 300,030 | 50003/16 | 50,003 | g4 |

Figure 3: Editable sample table. Rows can be deselected, edited (e.g. the grouping variable), and manually reordered; the generated queue reflects the edits.

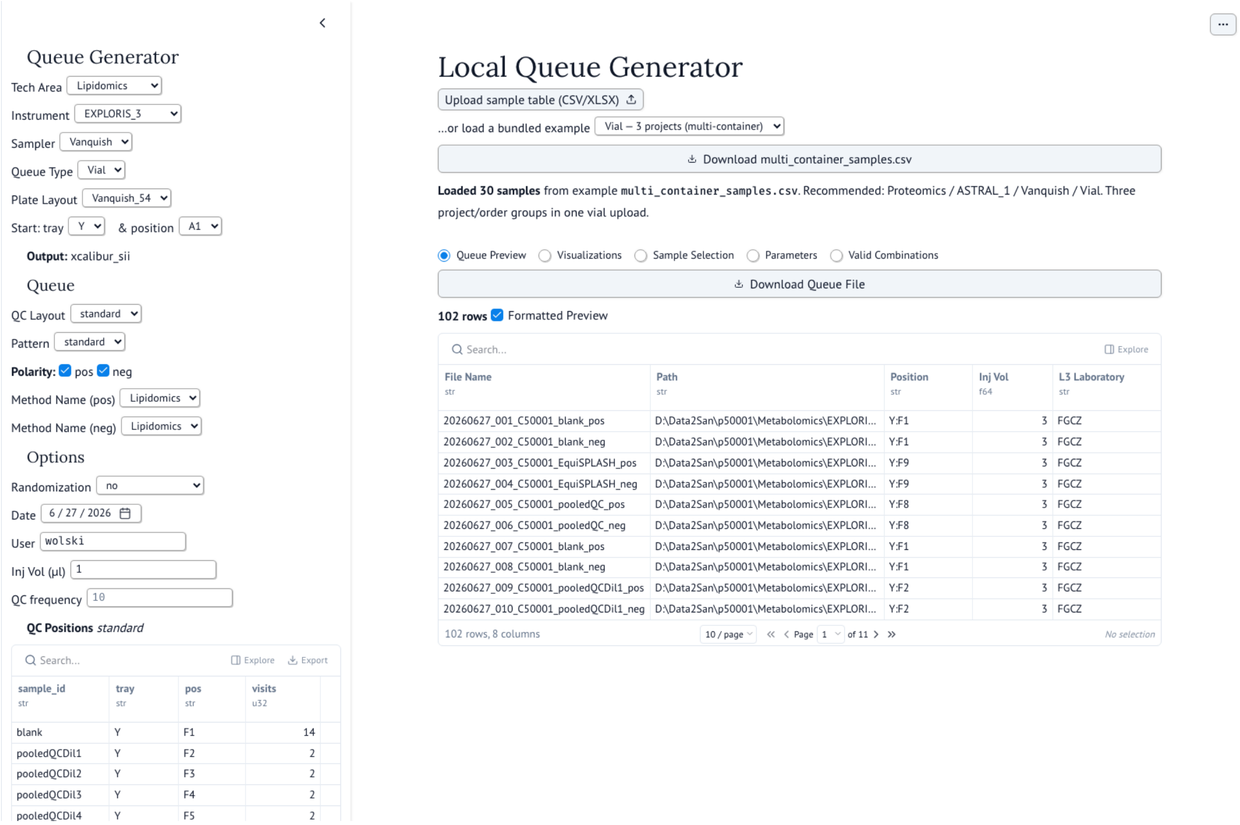


| sample_id | tray | pos | visits |
|---|---|---|---|
| blank | Y | F1 | 14 |
| pooledQCDil1 | Y | F2 | 2 |
| pooledQCDil2 | Y | F3 | 2 |
| pooledQCDil3 | Y | F4 | 2 |
| pooledQCDil4 | Y | F5 | 2 |

| File Name | Path | Position | Inj Vol | L3 Laboratory |
|---|---|---|---|---|
| 20260627_001_C50001_blank_pos | D:\Data2San\p50001\Metabolomics\EXPLORI... | Y:F1 | 3 | FGCZ |
| 20260627_002_C50001_blank_neg | D:\Data2San\p50001\Metabolomics\EXPLORI... | Y:F1 | 3 | FGCZ |
| 20260627_003_C50001_EquiSPLASH_pos | D:\Data2San\p50001\Metabolomics\EXPLORI... | Y:F9 | 3 | FGCZ |
| 20260627_004_C50001_EquiSPLASH_neg | D:\Data2San\p50001\Metabolomics\EXPLORI... | Y:F9 | 3 | FGCZ |
| 20260627_005_C50001_pooledQC_pos | D:\Data2San\p50001\Metabolomics\EXPLORI... | Y:F8 | 3 | FGCZ |
| 20260627_006_C50001_pooledQC_neg | D:\Data2San\p50001\Metabolomics\EXPLORI... | Y:F8 | 3 | FGCZ |
| 20260627_007_C50001_blank_pos | D:\Data2San\p50001\Metabolomics\EXPLORI... | Y:F1 | 3 | FGCZ |
| 20260627_008_C50001_blank_neg | D:\Data2San\p50001\Metabolomics\EXPLORI... | Y:F1 | 3 | FGCZ |
| 20260627_009_C50001_pooledQCDil1_pos | D:\Data2San\p50001\Metabolomics\EXPLORI... | Y:F2 | 3 | FGCZ |
| 20260627_010_C50001_pooledQCDil1_neg | D:\Data2San\p50001\Metabolomics\EXPLORI... | Y:F2 | 3 | FGCZ |

Figure 4: Formatted queue preview. The previewed table is the exact vendor file written to disk; the output format is a parameter, so the same queue can be emitted for any supported instrument.

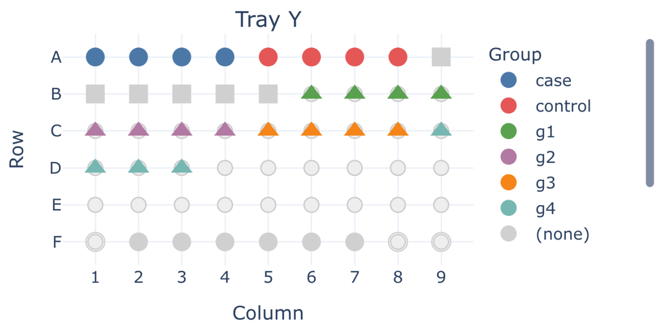


Figure 5: Plate-layout visualization coloured by grouping variable, annotated with a correlation-ratio ($\eta^2$) balance score for group versus plate position.

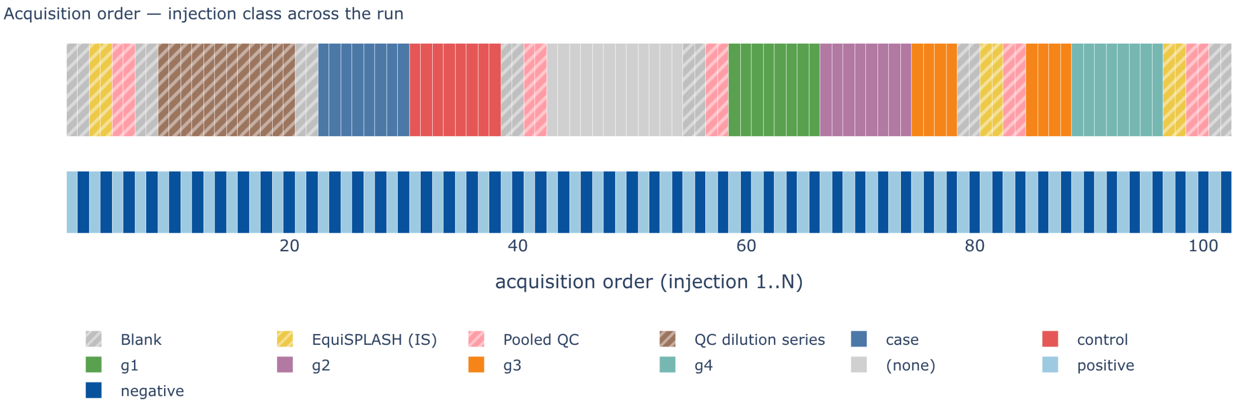


Figure 6: Acquisition-timeline visualization: the injection sequence coloured by injection class (biological group for user samples, QC type for QC injections) — or, alternatively, by QC cadence — annotated with the group versus run-position balance score.

#### 2 Configuration-editor walkthrough

The configuration editor exposes each configuration domain as an editable, validated table. Its overview tab (Figure 7) resolves a chosen technology/instrument/sampler/layout selection into the patterns, instrument configs, QC layouts, and methods it implies; the per-domain tabs that follow are representative. Every change is checked by the same cross-domain validation that runs at load before it is written.

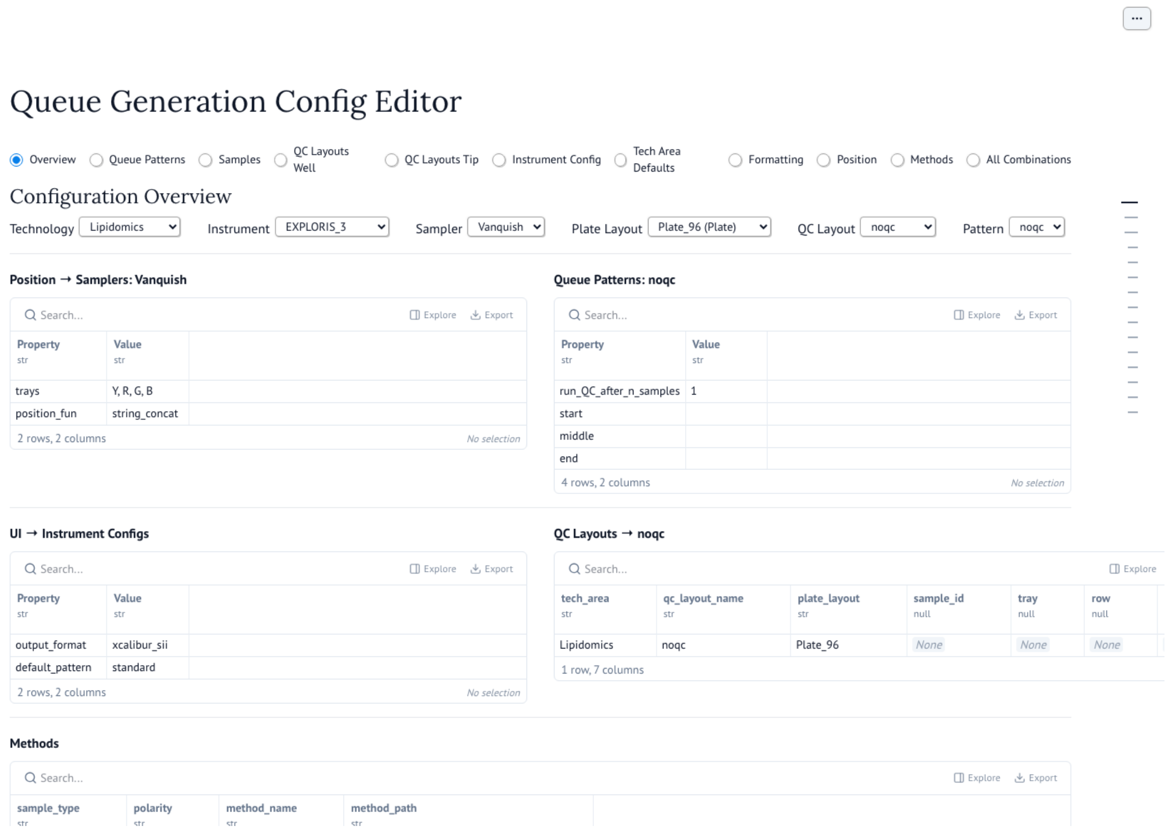


Figure 7: Configuration-editor overview tab. Tabs across the top expose each configuration domain — queue patterns, samples, QC layouts, instrument config, formatting, position, and methods — as editable tables; the overview resolves a chosen technology/instrument/sampler/layout selection into the patterns, instrument configs, QC layouts, and methods it implies. `Validate` re-runs the full cross-domain validation before changes are written, and `Submit for Review` records them as a reviewed commit.

QC-injection patterns are edited as `start/middle/end/separation` blocks with a recurrence count (Figure 8); the QC and standard samples those blocks reference are defined per technology (Figure 9).

Instruments, their UI defaults, and their acquisition methods are configured in separate domains (Figures 10–11), while reserved QC positions are declared per sampler/layout (Figure 12).

Finally, the editor surfaces the cross-domain validity of every instrument, sampler, layout, and pattern combination (Figure 13), so a misconfiguration is visible before any queue is generated.

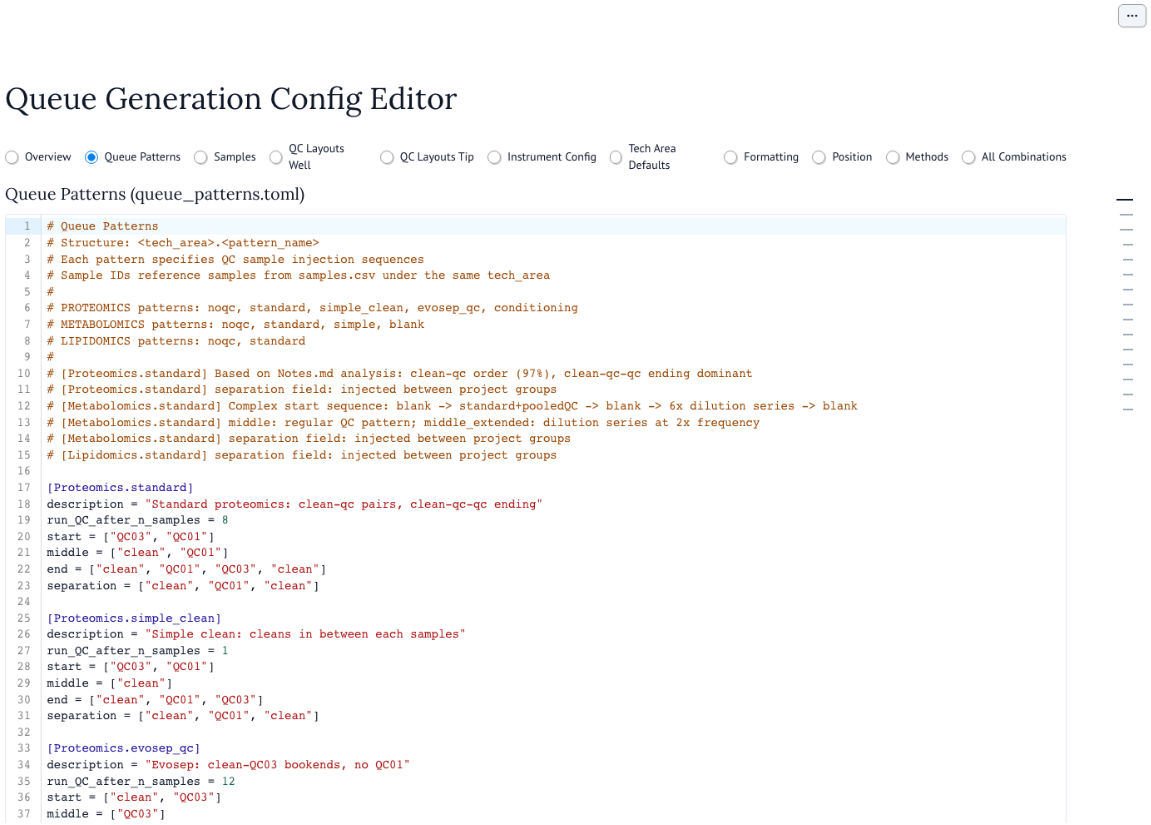


Figure 8: Queue-patterns editor: the QC-injection blocks and recurrence for each technology/pattern.

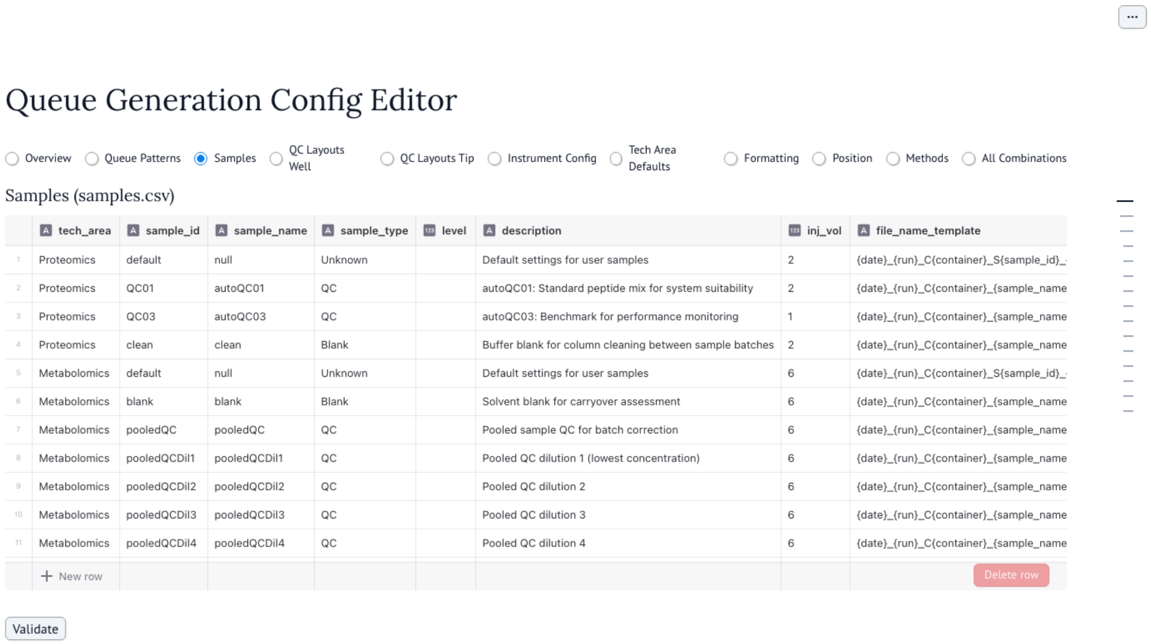

### Queue Generation Config Editor

Overview | Queue Patterns | Samples | QC Layouts Well | QC Layouts Tip | Instrument Config | Tech Area Defaults | Formatting | Position | Methods | All Combinations

Samples (samples.csv)

| | tech_area | sample_id | sample_name | sample_type | level | description | inj_vol | file_name_template |
|---|---|---|---|---|---|---|---|---|
| 1 | Proteomics | default | null | Unknown | | Default settings for user samples | 2 | {date}_{run}_C{container}_S{sample_id}_ |
| 2 | Proteomics | QC01 | autoQC01 | QC | | autoQC01: Standard peptide mix for system suitability | 2 | {date}_{run}_C{container}_{sample_name |
| 3 | Proteomics | QC03 | autoQC03 | QC | | autoQC03: Benchmark for performance monitoring | 1 | {date}_{run}_C{container}_{sample_name |
| 4 | Proteomics | clean | clean | Blank | | Buffer blank for column cleaning between sample batches | 2 | {date}_{run}_C{container}_{sample_name |
| 5 | Metabolomics | default | null | Unknown | | Default settings for user samples | 6 | {date}_{run}_C{container}_S{sample_id}_ |
| 6 | Metabolomics | blank | blank | Blank | | Solvent blank for carryover assessment | 6 | {date}_{run}_C{container}_{sample_name |
| 7 | Metabolomics | pooledQC | pooledQC | QC | | Pooled sample QC for batch correction | 6 | {date}_{run}_C{container}_{sample_name |
| 8 | Metabolomics | pooledQCDil1 | pooledQCDil1 | QC | | Pooled QC dilution 1 (lowest concentration) | 6 | {date}_{run}_C{container}_{sample_name |
| 9 | Metabolomics | pooledQCDil2 | pooledQCDil2 | QC | | Pooled QC dilution 2 | 6 | {date}_{run}_C{container}_{sample_name |
| 10 | Metabolomics | pooledQCDil3 | pooledQCDil3 | QC | | Pooled QC dilution 3 | 6 | {date}_{run}_C{container}_{sample_name |
| 11 | Metabolomics | pooledQCDil4 | pooledQCDil4 | QC | | Pooled QC dilution 4 | 6 | {date}_{run}_C{container}_{sample_name |



Figure 9: Samples editor: the per-technology QC and standard-injection catalogue (injection volume, sample type, file-name template) that patterns and QC layouts reference.

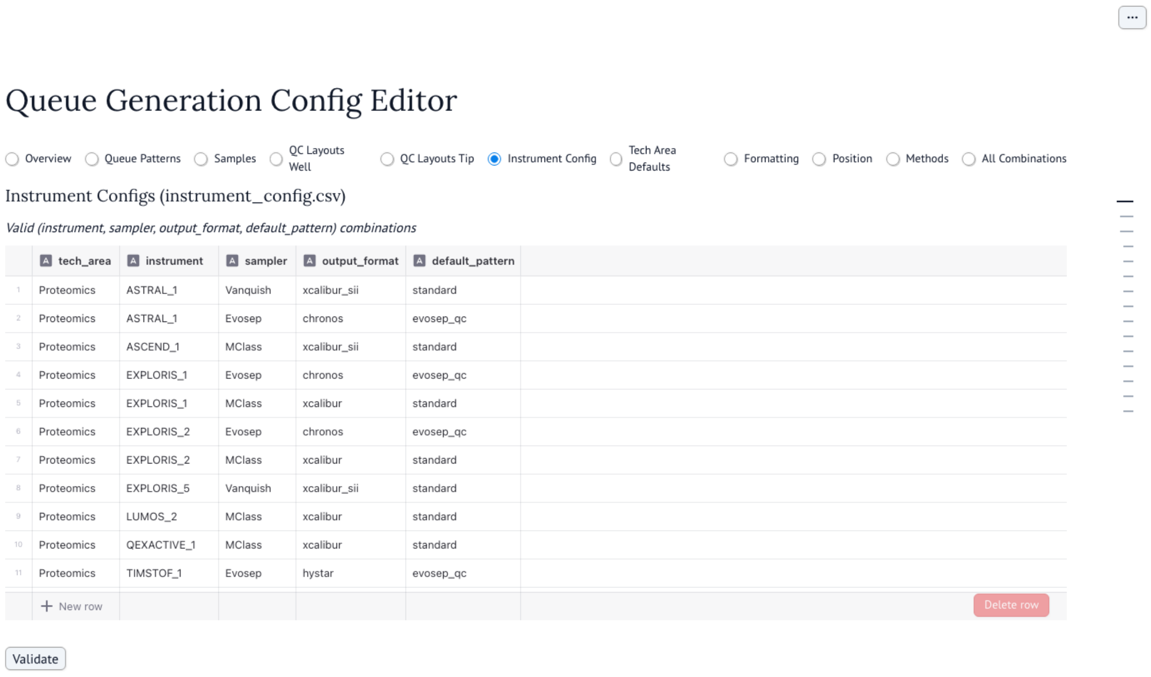


| | tech_area | instrument | sampler | output_format | default_pattern |
|---|---|---|---|---|---|
| 1 | Proteomics | ASTRAL_1 | Vanquish | xcalibur_sii | standard |
| 2 | Proteomics | ASTRAL_1 | Evosep | chronos | evosep_qc |
| 3 | Proteomics | ASCEND_1 | MClass | xcalibur_sii | standard |
| 4 | Proteomics | EXPLORIS_1 | Evosep | chronos | evosep_qc |
| 5 | Proteomics | EXPLORIS_1 | MClass | xcalibur | standard |
| 6 | Proteomics | EXPLORIS_2 | Evosep | chronos | evosep_qc |
| 7 | Proteomics | EXPLORIS_2 | MClass | xcalibur | standard |
| 8 | Proteomics | EXPLORIS_5 | Vanquish | xcalibur_sii | standard |
| 9 | Proteomics | LUMOS_2 | MClass | xcalibur | standard |
| 10 | Proteomics | QEXACTIVE_1 | MClass | xcalibur | standard |
| 11 | Proteomics | TIMSTOF_1 | Evosep | hystar | evosep_qc |

Figure 10: Instrument-config editor: the per-instrument UI defaults (sampler, output format, default pattern) that populate the queue-app menus.

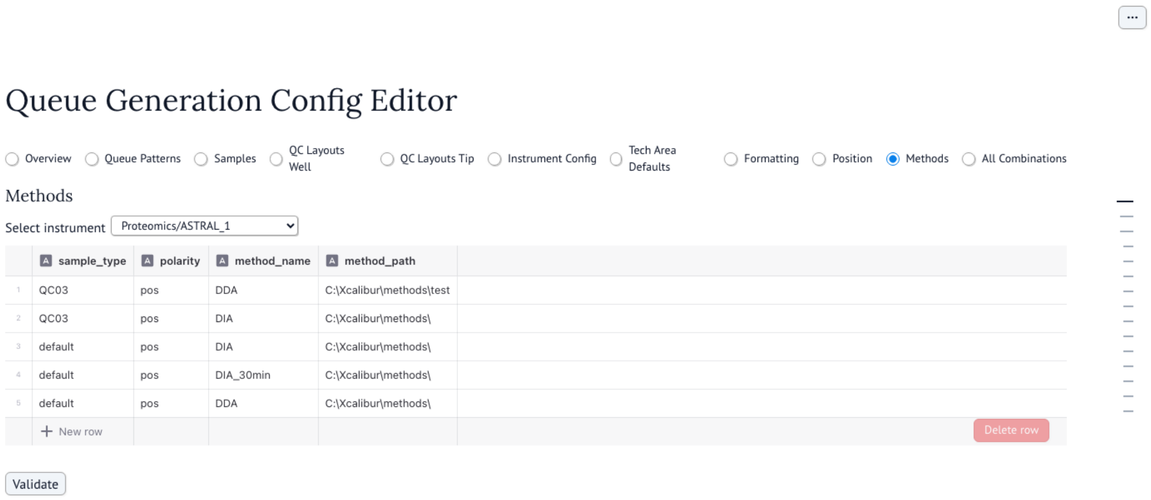


Figure 11: Methods editor: per-instrument acquisition methods by sample type and polarity, with their on-instrument paths.

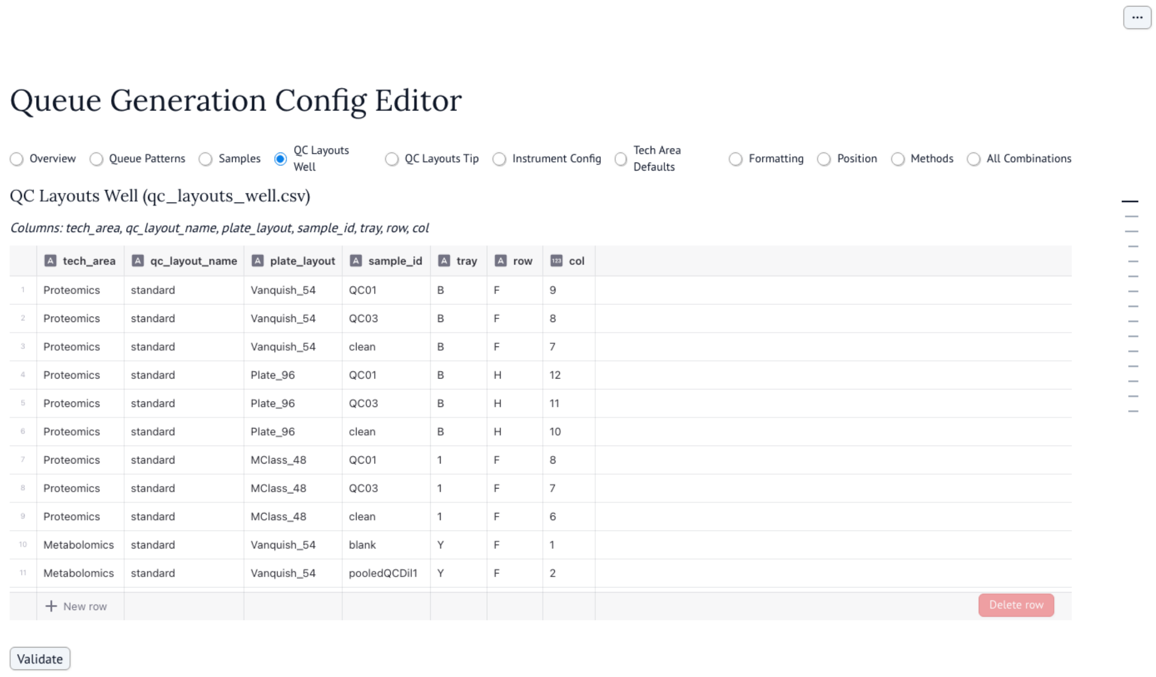


Figure 12: QC-layout editor (well-plate samplers): the reserved well positions QC injections always map to, independent of user-sample ordering.

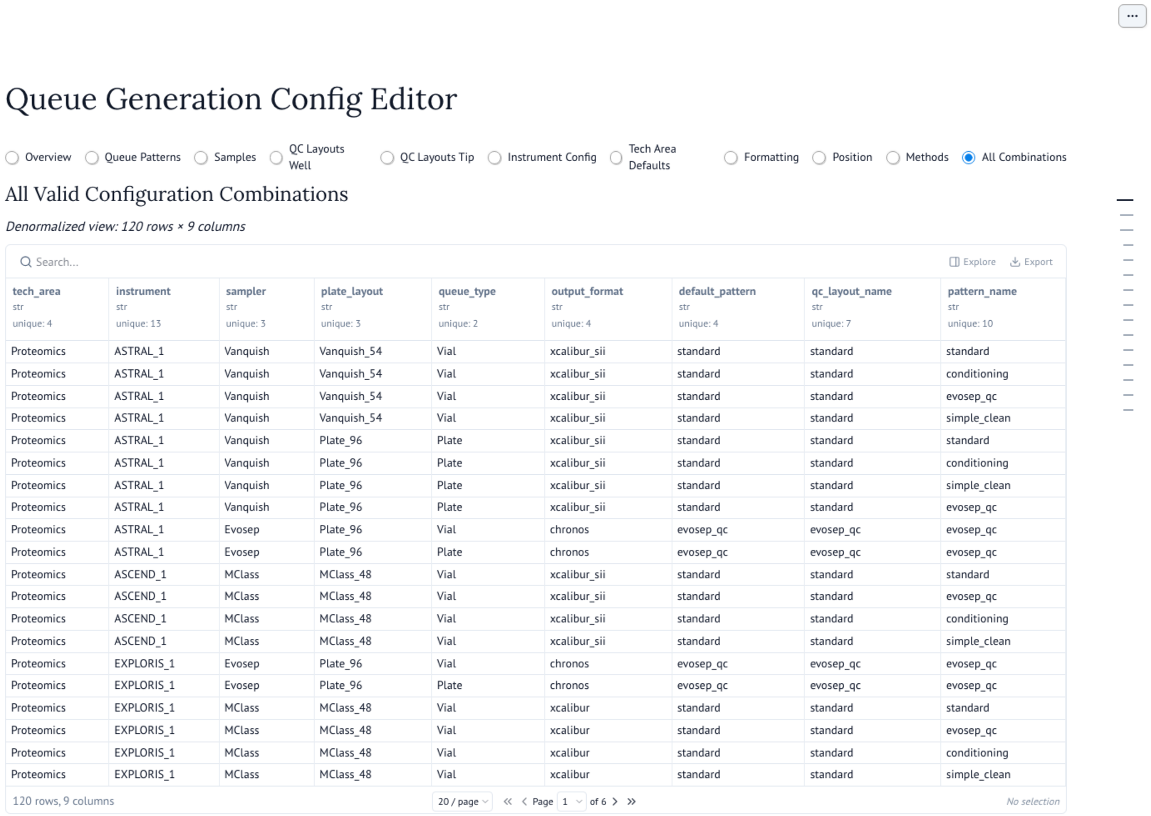


| tech_area<br>str<br>unique: 4 | instrument<br>str<br>unique: 13 | sampler<br>str<br>unique: 3 | plate_layout<br>str<br>unique: 3 | queue_type<br>str<br>unique: 2 | output_format<br>str<br>unique: 4 | default_pattern<br>str<br>unique: 4 | qc_layout_name<br>str<br>unique: 7 | pattern_name<br>str<br>unique: 10 |
|---|---|---|---|---|---|---|---|---|
| Proteomics | ASTRAL_1 | Vanquish | Vanquish_54 | Vial | xcalibur_sii | standard | standard | standard |
| Proteomics | ASTRAL_1 | Vanquish | Vanquish_54 | Vial | xcalibur_sii | standard | standard | conditioning |
| Proteomics | ASTRAL_1 | Vanquish | Vanquish_54 | Vial | xcalibur_sii | standard | standard | evosep_qc |
| Proteomics | ASTRAL_1 | Vanquish | Vanquish_54 | Vial | xcalibur_sii | standard | standard | simple_clean |
| Proteomics | ASTRAL_1 | Vanquish | Plate_96 | Plate | xcalibur_sii | standard | standard | standard |
| Proteomics | ASTRAL_1 | Vanquish | Plate_96 | Plate | xcalibur_sii | standard | standard | conditioning |
| Proteomics | ASTRAL_1 | Vanquish | Plate_96 | Plate | xcalibur_sii | standard | standard | simple_clean |
| Proteomics | ASTRAL_1 | Vanquish | Plate_96 | Plate | xcalibur_sii | standard | standard | evosep_qc |
| Proteomics | ASTRAL_1 | Evosep | Plate_96 | Vial | chronos | evosep_qc | evosep_qc | evosep_qc |
| Proteomics | ASTRAL_1 | Evosep | Plate_96 | Plate | chronos | evosep_qc | evosep_qc | evosep_qc |
| Proteomics | ASCEND_1 | MClass | MClass_48 | Vial | xcalibur_sii | standard | standard | standard |
| Proteomics | ASCEND_1 | MClass | MClass_48 | Vial | xcalibur_sii | standard | standard | evosep_qc |
| Proteomics | ASCEND_1 | MClass | MClass_48 | Vial | xcalibur_sii | standard | standard | conditioning |
| Proteomics | ASCEND_1 | MClass | MClass_48 | Vial | xcalibur_sii | standard | standard | simple_clean |
| Proteomics | EXPLORIS_1 | Evosep | Plate_96 | Vial | chronos | evosep_qc | evosep_qc | evosep_qc |
| Proteomics | EXPLORIS_1 | Evosep | Plate_96 | Plate | chronos | evosep_qc | evosep_qc | evosep_qc |
| Proteomics | EXPLORIS_1 | MClass | MClass_48 | Vial | xcalibur | standard | standard | standard |
| Proteomics | EXPLORIS_1 | MClass | MClass_48 | Vial | xcalibur | standard | standard | evosep_qc |
| Proteomics | EXPLORIS_1 | MClass | MClass_48 | Vial | xcalibur | standard | standard | conditioning |
| Proteomics | EXPLORIS_1 | MClass | MClass_48 | Vial | xcalibur | standard | standard | simple_clean |

Figure 13: All-combinations view: the resolved validity of each combination of technology, instrument, sampler, layout, and pattern across the configuration.

Table 1: Normalised sample-table columns accepted by the standalone upload, and equivalently derived from B-Fabric. A submission is read as *plate* when both `plate_id` and `grid_position` are present, otherwise as *vial*. Integer columns are validated on load.

| Canonical column | Role | Accepted aliases |
|---|---|---|
| sample_name | required | Sample Name, name |
| sample_id | required (int) | Sample ID, id |
| container_id | required (int) | Container ID |
| grouping_var | optional | grouping var, groupingvar_name |
| tube_id | optional (vial) | Tube ID, tubeid |
| plate_id | required for plate (int) | Plate ID |
| grid_position | required for plate | grid position, _gridposition |
| tray | optional (plate) | — |

### 3 Sample-table upload schema

In standalone (no-LIMS) mode `qg` ingests a CSV or XLSX sample table. The parser normalises common header spellings — internal snake_case names, exported display names, and the B-Fabric export field names — to the canonical columns in Table 1, and reads the submission as a *plate* table when both `plate_id` and `grid_position` are present, otherwise as a *vial* table. This is the same normalised schema `qg` derives from a B-Fabric order, so a table exported from the LIMS and a hand-prepared spreadsheet drive the generator identically.

### 4 A worked reproducibility example

Every randomized run records the seed it used, and the exported parameters are *self-contained*: besides the seed and the full sample list, `qg` embeds its own version and a `resolved_config` snapshot of exactly the configuration the run drew on — the one instrument, sampler, pattern, plate and QC layouts, output format, and the referenced sample and method definitions. The block below is taken verbatim from a generated queue (a 12-sample, three-group proteomics run with `blocked_uniform` randomization); the operator left `seed` unset, so `qg` drew one and wrote it back:

```
"qg_version": "0.8.0",
"parameters": {
  "tech_area": "Proteomics", "instrument": "ASTRAL_1",
  "sampler": "Vanquish", "output_format": "xcalibur_sii",
  "queue_pattern": "standard", "queue_type": "Vial",
  "plate_layout": "Vanquish_54", "qc_layout_name": "standard",
  "polarity": ["pos"], "randomization": "blocked_uniform",
  "seed": 1883415883
},
"resolved_config": {       // the configuration this run used, inlined
  "instrument": {...}, "queue_pattern": {...}, "plate_layout": {...},
  "sampler": {...}, "samples": [...], "methods": [...]
}
```

The exported JSON also carries the full sample list. Re-running the command-line interface on that document reproduces the queue byte-for-byte, without the GUI and without the external configuration:

```
uv run qg params.json -o queue.csv
```

Because the file carries the seed, the `qg` version, and the resolved configuration, the same document yields the same queue on any machine — independent of the local `qg_configs` tree and immune to

Table 2: Instruments shipped per technology (15 total), across three autosamplers (well-plate Vanquish and M-Class; tip-based Evosep), three plate layouts, and four vendor output formats resolving to three writers (XCalibur CSV, Chronos CSV, HyStar XML).

| Technology | Instruments |
|---|---|
| Proteomics | ASTRAL_1, ASCEND_1, EXPLORIS_1, EXPLORIS_2, EXPLORIS_5, LUMOS_2, QEXACTIVE_1, [illegible] |
| Metabolomics | EXPLORIS_3, QEXACTIVEHF_2, QUANTIVA_1 |
| Lipidomics | EXPLORIS_3, EXPLORIS_4, QEXACTIVEHF_2 |

Table 3: QC-injection patterns shipped per technology (11 total). Each is a declarative `start/middle/end/separation` block set with a `run_QC_after_n_samples` recurrence, selectable per queue.

| Technology | QC patterns |
|---|---|
| Proteomics | standard, simple_clean, evosep_qc, conditioning |
| Metabolomics | standard, simple, blank, cal_series, noqc |
| Lipidomics | standard, noqc |

later edits of it; only the deterministic modes (`no`) are seed-independent. This exact run ships as a bundled example (`qg.examples.params/repro_proteomics_12.json`), runnable directly with `qg repro_proteomics_12.json`.

### 5 Shipped configuration

The configuration distributed with `qg` spans a realistic core-facility instrument park (Table 2) and a set of QC-injection patterns per technology (Table 3); an internal test-fixture technology is excluded from both. All are configuration entries, not code, and are the extension points for new instruments, samplers, layouts, patterns, and formats.

### 6 Configuration examples

The two longer configuration blocks referenced in the main text are reproduced here in full. The metabolomics `standard` QC pattern brackets the run with solvent blanks (`blank`), amino-acid and organic-acid/phosphate standard mixes (`108mix_AA`, `108mix_OAP`), pooled-sample QC (`pooledQC`), and a six-step pooled-QC dilution series (`pooledQCDil1`–`pooledQCDil6`), and re-injects that series every second `middle` block through the optional `middle_extended` field (the long arrays are wrapped here for column width):

```
[Metabolomics.standard]
run_QC_after_n_samples = 10
start = [
  "blank", "108mix_AA", "108mix_OAP",
  "pooledQC", "blank",
  "pooledQCDil1", "pooledQCDil2",
  "pooledQCDil3", "pooledQCDil4",
  "pooledQCDil5", "pooledQCDil6",
  "pooledQC", "blank",
]
middle = [
  "pooledQC", "108mix_AA",
  "108mix_OAP", "blank",
]
end = [
  "pooledQC", "108mix_AA",
  "108mix_OAP", "blank",
]
separation = ["blank", "pooledQC"]
middle_extended = [
  "pooledQCDil1", "pooledQCDil2",
  "pooledQCDil3", "pooledQCDil4",
  "pooledQCDil5", "pooledQCDil6",
]
middle_extended_frequency_multiplier = 2
```

Each output format is a declarative column map in `output_formats.toml`, mapping internal queue fields to the vendor's worklist columns. The XCalibur SII map sets eight base columns and lets metabolomics add two more, inheriting the rest:

```
[xcalibur_sii.columns]
"File Name"         = "file_name"
Path                = "data_path"
Position            = "position"
"Inj Vol"           = "inj_vol"
"L3 Laboratory"     = "literal:FGCZ"
"Sample ID"         = "sample_id"
"Sample Name"       = "sample_name"
"Instrument Method" = "method"

# Metabolomics inherits, then adds:
[xcalibur_sii.columns.Metabolomics]
"Sample Type" = "sample_type"
"Level"       = "level"
```

File and folder names are generated from templates. Each `instruments.csv` row carries a `path_template` built from `{container}`, `{user}`, and `{date}`, and each `samples.csv` row a `file_name_template` restricted to a fixed placeholder set (`{date}`, `{run}`, `{container}`, `{sample_id}`, `{sample_name}`, `{polarity}`, `{method_name}`, `{level}`, `{concentration}`). The example below shows the Proteomics `ASTRAL_1` path template and the default-sample file-name template, with the values one queue substitutes into them:

```
# instruments.csv  path_template  (Proteomics, ASTRAL_1)
D:\Data2San\p{container}\Proteomics\ASTRAL_1\{user}_{date}

# samples.csv  file_name_template  (Proteomics, default sample)
{date}_{run}_C{container}_S{sample_id}_{sample_name}

# rendered for user=cpanse, date=20260112, container=37180,
#          run=007, sample_id=123456, sample_name=HeLa_A1
data path:  D:\Data2San\p37180\Proteomics\ASTRAL_1\cpanse_20260112
file name:  20260112_007_C37180_S123456_HeLa_A1
```

### 7 Randomization quality across seeds and imbalance

The main-text randomization figure (Figure 4) reports a single seed on one unbalanced design. The two figures here generalize that result with the real `qg` randomization engine. Figure 14 repeats all four modes over 2000 independent seeds and adds a second, clustering-sensitive metric (the longest consecutive run of one biological group), and Figure 15 sweeps the degree of group imbalance.

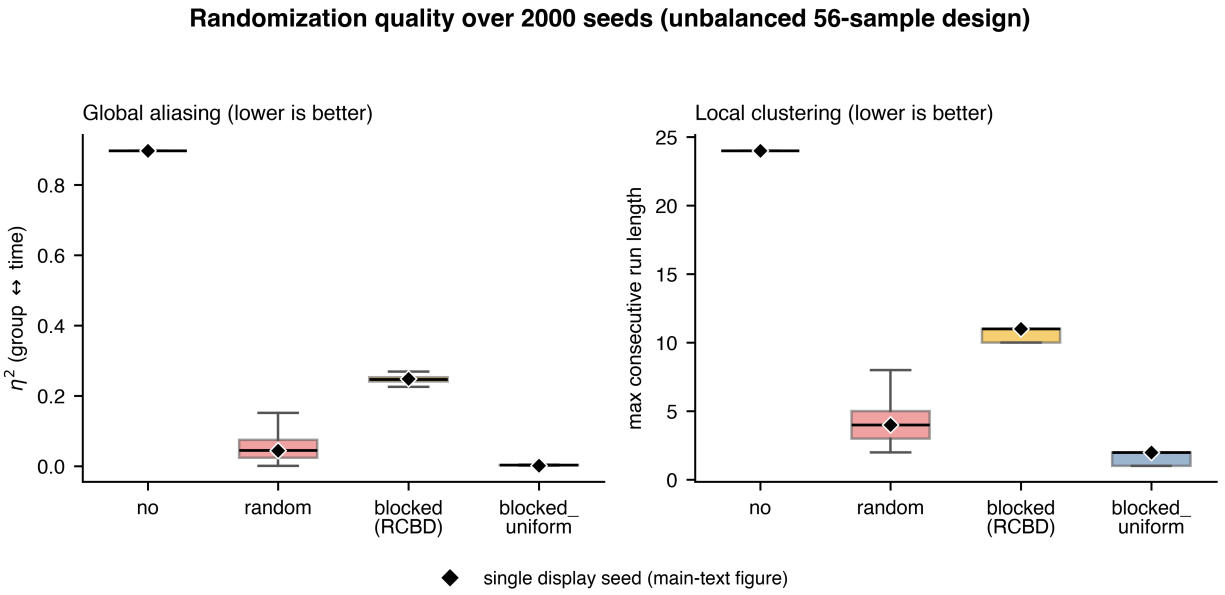


Figure 14: Randomization quality over 2000 independent seeds on the unbalanced 56-sample design ($A = 24$, $B = 14$, $C = 10$, $D = 8$) of the main text. **Left:** the distribution of $\eta^2$ (the share of acquisition-position variance explained by biological group; lower is better) for each mode. **Right:** the longest consecutive run of one biological group (local clustering; lower is better). Boxes span the interquartile range; whiskers the range excluding outliers; the diamond marks the single display seed used in the main-text figure. `blocked_uniform` dominates on both metrics (median $\eta^2 \approx 0.003$, run length $\approx 2$); `random` decorrelates group from time on average but leaves local clustering (run length $\approx 4$); and `blocked` (RCBD) is worse than `random` on $\eta^2$ (median $\approx 0.25$) for these unbalanced groups. Generated by `make_seed_sweep_figure.py`.

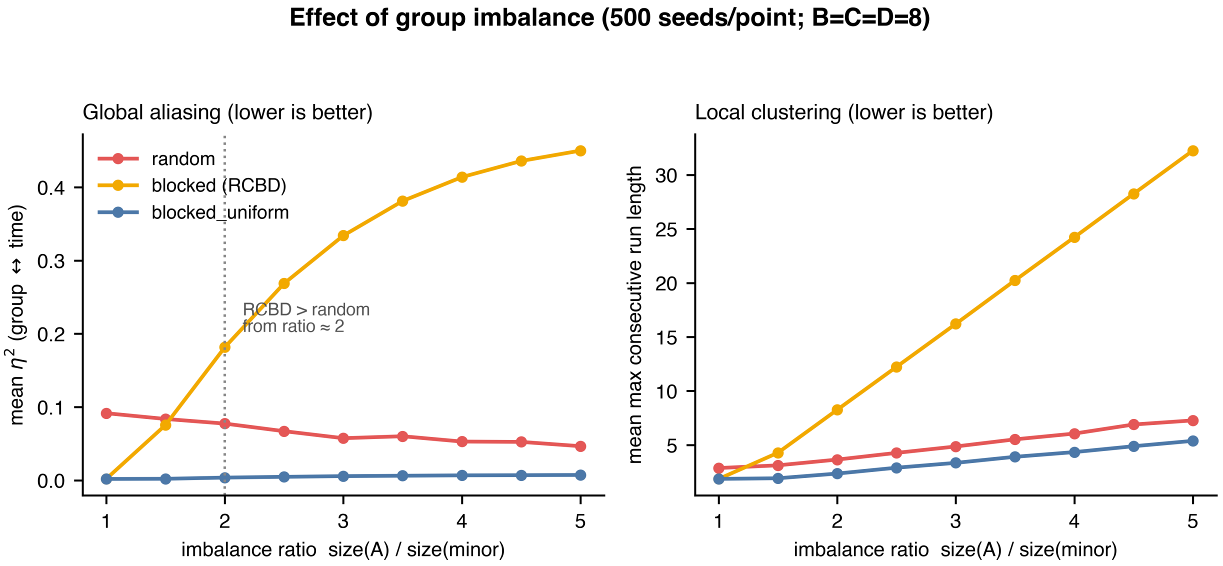


Figure 15: Effect of group imbalance. Three minor groups are held at 8 samples each while the dominant group grows from 8 (ratio 1, balanced) to 40 (ratio 5); 500 seeds per point. **Left:** mean $\eta^2$; **right:** mean longest run. On balanced groups RCBD is near-optimal, but its $\eta^2$ climbs past a plain `random` shuffle once the dominant group exceeds about twice the others (dotted line), reaching $\approx 0.45$ at ratio 5, whereas `blocked_uniform` stays near zero at every imbalance. Generated by `make_imbalance_sweep_figure.py`.

### 8 A worked lipidomics acquisition queue

Figure 16 shows a lipidomics queue generated by `qg` from the shipped `Lipidomics.standard` pattern, illustrating the lipidomics workflow described in the main text. The exact run ships as a self-contained bundled example, and the figure is the app's own *Acquisition Timeline* view of it — not a separate plot. A reader reproduces it interactively by running `qg-app-local` (`make app-local`), loading `lipidomics_standard.json` from the bundled-run dropdown (the app's reproduce mode), and opening *Visualizations* → *Acquisition Timeline*; the command line equivalent, `qg lipidomics_standard.json`, regenerates the identical queue from that one file.

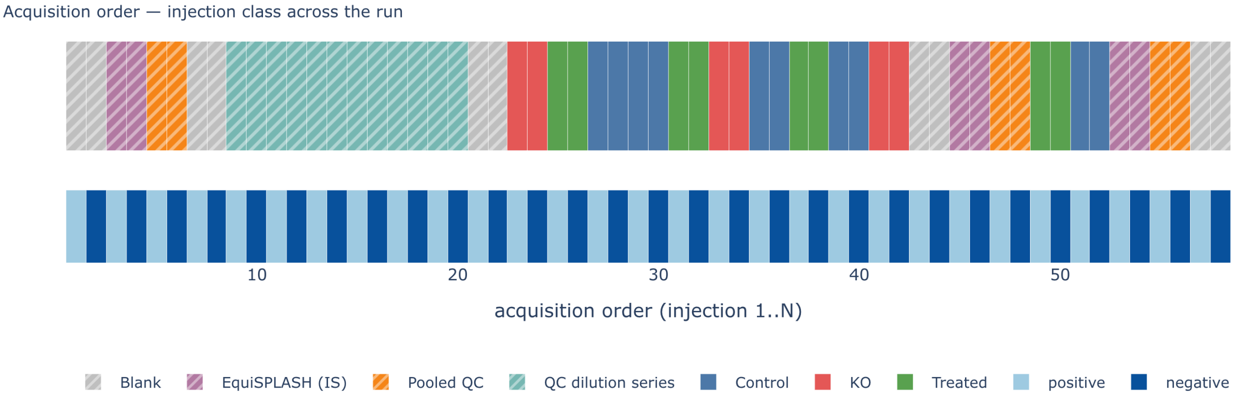


Figure 16: A worked lipidomics acquisition queue generated by *qg* from the shipped `Lipidomics.standard` pattern (EXPLORIS instrument, Vanquish sampler, dual-polarity acquisition, `blocked_uniform` run order). Twelve samples in three groups expand to 58 injections. The upper track colours each injection by class: a leading system-suitability block (solvent blanks, the EquiSPLASH class-spanning internal-standard mix, a pooled QC, and a six-point pooled-QC dilution series), then the user samples group-interleaved, with pooled QC and internal standards recurring through the run. The lower track shows that every injection — QC and sample alike — is acquired in both positive and negative ionization polarity. This is the app's *Acquisition Timeline* view, captured by `shot_viz.py` after loading the bundled `lipidomics_standard.json` in the app's reproduce mode.

### 9 Figure generation and computational reproducibility

Every figure in this Supporting Information and in the main text is produced programmatically by a committed script in the public `qg` repository under `docs/examples/figures/` (with a README mapping each script to its figure and giving run instructions), so a reader with the released tool can reproduce every one; nothing is hand-drawn except the architecture and configuration schematics.

The analytical figures are rendered directly from the `qg` engine:

- the main-text randomization comparison — `make_randomization_figure.py`;
- Figure
  reffig:si-seedsweep, the 2000-seed sweep — `make_seed_sweep_figure.py`;
- Figure
  reffig:si-imbalance, the imbalance sweep — `make_imbalance_sweep_figure.py`;
- the main-text plate and acquisition-timeline panels — `make_plate_figure.py`, through the same `qg.viz` helpers the graphical interface uses.

The remaining figures are scripted captures of `qg`'s own graphical interface, taken headlessly from the standalone local app (`qg-app-local`) with playwright; the capture scripts (`shot_*.py`) ship alongside the figure scripts. In particular, the worked lipidomics queue (Figure 16) is the app's *Acquisition Timeline* view, captured by `shot_viz.py` after loading the bundled, self-contained `lipidomics_standard.json` in the app's reproduce mode — so the figure and the one-command example (`qg lipidomics_standard.json`) are the same run and cannot diverge. The plate-layout and acquisition-timeline interface panels (Figures 5, 6) are captured the same way from the bundled multi-container example. Only the architecture and configuration schematics are hand-authored diagrams.
